# A Safe and Effective Mucosal RSV Vaccine Consisting of RSV Phosphoprotein and Flagellin Variant

**DOI:** 10.1101/2021.02.09.430425

**Authors:** Bali Zhao, Jingyi Yang, Bing He, Xian Li, Hu Yan, Shuning Liu, Yi Yang, Dihan Zhou, Bowen Liu, Xuxu Fan, Maohua Zhong, Ejuan Zhang, Fan Zhang, Yue Zhang, Yao-Qing Chen, Shibo Jiang, Huimin Yan

## Abstract

Respiratory syncytial virus (RSV) is a major cause of serious acute lower respiratory tract infection in infants and the elderly. No licensed RSV vaccine available thus far calls for the development of vaccines with new target(s) and vaccination strategies. Here, we constructed a recombinant protein, designated P-KFD1, comprised of RSV phosphoprotein (P) and *E. coli* K12 strain-derived flagellin variant KFD1. Intranasal (i.n.) immunization with P-KFD1 inhibits RSV replication in both upper and lower respiratory tract, and protects mice against lung disease without vaccine-enhanced disease (VED). The P-specific CD4^+^ T cells provoked by P-KFD1 i.n. immunization, either reside in or migrate to respiratory tract, mediate protection against RSV infection. Sc-RNA seq and carboxyfluorescein succinimidyl ester (CFSE) labeled cell transfer further characterized the Th1 and Th17 responses induced by P-KFD1. Finally, we found the anti-viral protection depends on either IFN-γ or IL-17A. Collectively, P-KFD1 is promising as a safe and effective mucosal vaccine candidate to prevent RSV infection.

**HIGHLIGHTS:**

- A new subunit RSV vaccine candidate with new target and vaccination strategy, P-KFD1, is designed and generated
- Intranasal immunization with P-KFD1 protects mice against RSV infection and averts vaccine-enhanced disease
- Sc-RNA seq and CFSE-labelled cell transfer identified characteristics of the protective CD4^+^ T cells
- Local and peripheral CD4^+^ T cells provide protection against RSV infection dependent on either IFN-γ or IL-17A

## INTRODUCTION

Human respiratory syncytial virus (RSV) was first isolated and identified as an important cause of bronchiolitis in infants as early as 1956 (Blount et al., 1956; Chanock and Finberg, 1957). It remains a major cause of severe acute lower respiratory tract infection (ALRI) not only in infants, but also in young children, the elderly, and immunocompromised populations worldwide (Chemaly et al., 2014; Falsey et al., 2005; Nair et al., 2010). RSV infection can result in severe complications, including bronchiolitis or pneumonia, often with acute respiratory distress, requiring hospitalization and creating a heavy medical cost and burden for treatment and prevention all over the world. In 2015 alone, RSV infection resulted in about 3.2 million episodes of hospitalization across the globe and 59,600 deaths in children younger than 5 years (Shi et al., 2017). A recent multi-site case-control study found that RSV had the greatest etiological fraction as high as 31.1% of all bacteria and virus pathogens that caused severe pneumonia requiring hospital admission in children from Africa and Asia (Group, 2019). Even worse, natural RSV infection does not elicit long-lasting immunity, and repeated infections occur throughout life (Glezen et al., 1986; Henderson et al., 1979). Only one licensed monoclonal antibody product (Palivizumab) is currently used to reduce the frequency of severe disease in the high-risk neonates (Group, 1998). The development of a safe and effective RSV vaccine has been elusive thus far.

The intramuscularly administered formalin-inactivated RSV (FI-RSV) vaccine primed for enhanced illness in infants on natural RSV infection, this phenomenon was replicable in animal models and named as vaccine-enhanced disease (VED) (Chin et al., 1969; Fulginiti et al., 1969; Kapikian et al., 1969; Kim et al., 1969; Waris et al., 1996). Intensive investigation showed that VED is usually involved in Th2-biased immune response and substantial binding antibodies (Ruckwardt et al., 2019). Subsequent studies then adopted subunit-based vaccines, mostly by targeting cell surface fusion (F) protein or attachment glycoprotein (G), owing to their immunogenicity for inducing neutralizing antibodies (Graham et al., 2015). However, G protein-based vaccines also primed for immunopathology in animals (Johnson et al., 1998; Openshaw et al., 1992; Tebbey et al., 1998), making G-targeted vaccines much more complicated. The elucidation of RSV F protein pre-fusion (pre-F) conformation (McLellan et al., 2013b) and its potential as an immunogen to induce potent neutralizing antibodies (Crank et al., 2019; Marcandalli et al., 2019; McLellan et al., 2013a) gave new hope for the development of F protein-based RSV vaccines. However, some previous clinical studies of post-fusion F (post-F)-based vaccines have failed (Ruckwardt et al., 2019). Consequently, new RSV vaccine targets and vaccination strategies warrant intensive investigation.

Because RSV infection mostly initiates from the upper respiratory tract and is restricted to the lung, an RSV vaccine could prophylactically prevent infection at upper and lower respiratory tracts if mucosal immunity could be elicited effectively in the respiratory mucosa (Yang and Varga, 2014). Moreover, appropriate immunization routes, such as intranasal (i.n.) administration, and specific adjuvant formulations to induce potent and broad mucosal immune responses are needed. Therefore, we first chose the cholera toxin B subunit (CTB) as a mucosal adjuvant to test whether the RSV internal phosphoprotein (P), nucleoprotein (N), non-structural protein 1 (NS-1) and M2-1 could be potential vaccine targets in BALB/c mice. We found that i.n. administration of P, together with CTB (P+CTB), significantly reduced viral loads in both noses and lungs of mice upon RSV challenge compared with mock immunized mice. This encouraging result urged us to further evaluate P as a vaccine target by engaging more mucosal adjuvants and immunization strategies.

However, certain degree of increased inflammatory cell infiltration and mucus production were observed in the P+CTB immunized mice compared to saline immunized mice post RSV challenge, suggesting potential vaccine-induced immunopathogy. Thus, we further tried another mucosal adjuvant flagellin (Mizel and Bates, 2010), and integrated flagellin with the newly identified vaccine target P, as we did for the development of anti-caries vaccines (Bao et al., 2015; Sun et al., 2012; Yang et al., 2017). The resultant fusion protein-based vaccine P-KFD1 was comprised of phosphoprotein (P) of RSV and *E. coli* K12 strain-derived flagellin (KF) with deletion of its hypervariable domain (KFD1) (Donnelly and Steiner, 2002; Yang et al., 2013), where the P and KFD1 were covalently coupled. Here, we report that i.n. immunization with P-KFD1 protects mice against RSV infection, as well as lung disease.

## RESULTS

### P-specific immune responses induced by intranasal immunization with P-KFD1 or P+CTB protect mice against RSV infection

We generated a recombinant protein P-KFD1 (**Figures S1A and S1B**), and investigated the immunogenicity and anti-viral efficacy of P-KFD1 as a potential mucosal subunit RSV vaccine targeting on P. P-KFD1 retains Toll like receptor 5 (TLR5) activity at a marginal lower level than full length KF (**Figure S1C**). We firstly evaluated the immunogenicity of P by i.n. immunization with P-KFD1 or cholera toxin B subunit (CTB) adjuvanted P protein in BALB/c mice. Briefly, 20 μg of P-KFD1 in saline or 10 μg of P protein mixed with 2 μg of CTB (P+CTB) were prepared as one dose of immunogen. BALB/c mice were immunized with P-KFD1, P+CTB or saline only for sham control, respectively, at weeks (w) 0, 4 and 8 (**Figure 1A, upper panel**). P-specific IFN-γ secreting T cell response could be detected in both lungs and spleens of P-KFD1 or P+CTB immunized mice (**Figure 1B**). Meanwhile, high levels of P-specific saliva IgA, serum IgG and serum IgA antibody responses could be induced by i.n. immunization with either P-KFD1 or P+CTB compared to sham immunization (**Figure 1C**). These results indicated that recombinant P is immunogenic in mice by i.n. immunization in either P-KFD1 or P+CTB formulation, although the levels of P-specific immune responses induced by P-KFD1 were lower than P+CTB (**Figures 1B and 1C**). Next, the protective efficacy against RSV infection by i.n. immunization with P-KFD1 or P+CTB was evaluated and compared with that by intramuscular (i.m.) immunization of formalin-inactivated RSV (FI-RSV) in mice. Thus, one more group of mice was immunized with FI-RSV at weeks 8 and 10 with Alum as adjuvant (**Figure 1A, lower panel**). All immunized mice were challenged with RSV strain A2 at week 12, and sacrificed at 4 days post challenge for evaluating viral loads in noses and lungs (**Figure 1A**). Compared with control group, P-KFD1, P+CTB or FI-RSV immunized mice significantly reduced viral loads in both noses and lungs (**Figure 1D**). In addition, viral loads decrease could not be observed in either CTB only or KFD1 only intranasally immunized mice (**Figures 1E and 1F**), suggesting that i.n. immunization of P-KFD1 or P+CTB elicited P-specific immune responses conferred protection against RSV infection.

**Figure 1.**
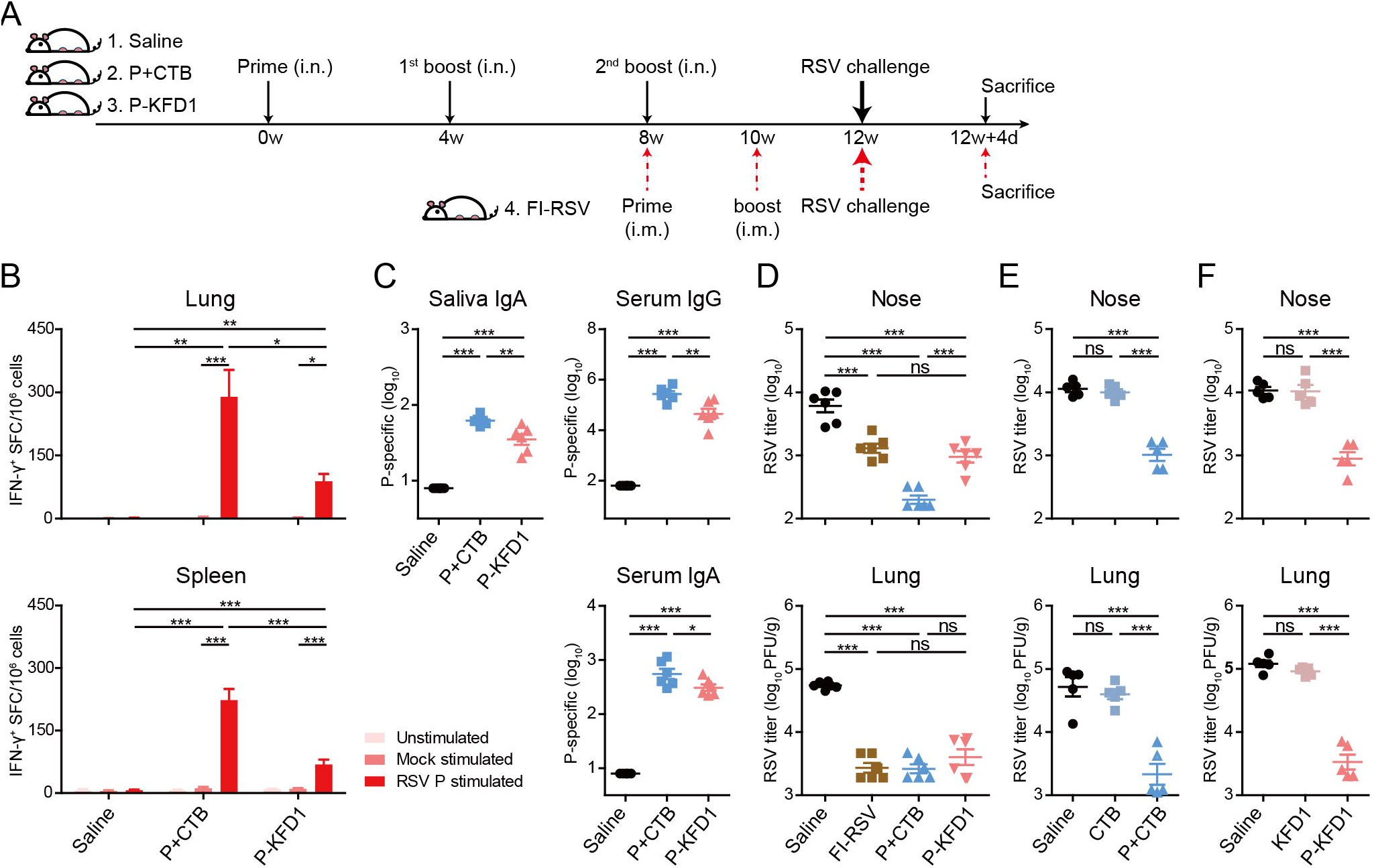
P-specific immune responses induced by intranasal immunization with P-KFD1 or P+CTB protect mice against RSV infection. (**A**) Immunization schedule. Groups of BALB/c mice were immunized with saline, P+CTB, P-KFD1 or FI-RSV at indicated time points, respectively, prior to infection with 2 ⅹ 10^6^ PFU of RSV A2 and sacrificed at 4 days post RSV infection (dpi). (**B**) P-specific IFN-γ secreting T cells were determined by ELISpot and were expressed as IFN-γ^+^ spot forming cells (SFC)/ million cells at 7 days post last immunization, n = 5 mice per group. (**C**) P-specific saliva IgA, serum IgG and serum IgA antibody titers in P+CTB or P-KFD1 immunized mice were determined by ELISA at 14 days post last immunization, n = 6 mice per group. (**D**) RSV titers in saline, FI-RSV, P+CTB or P-KFD1 immunized mice were monitored at 4 dpi, n = 6 mice per group. (**E**) RSV titers in saline, CTB or P+CTB immunized mice were measured at 4 dpi, n = 5 mice per group. (**F**) RSV titers in saline, KFD1 or P-KFD1 immunized mice were detected at 4 dpi, n = 5 mice per group. Data are represented as mean ± SEM of three independent experiments. In **Figure 1B**, two columns were compared using unpaired *t* test. In **Figures 1C to 1F**, groups were compared using one-way ANOVA. * p < 0.05, ** p < 0.01, *** p < 0.001, ns means non-significant. See also **Figure S1**.

### Intranasal immunization with P-KFD1 protects mice against lung disease caused by RSV infection

A primary concern for development of an RSV vaccine is the potential for VED that occurred in FI-RSV vaccinated infants upon natural RSV infection. We thus evaluated the pathological changes in P-KFD1 or P+CTB immunized mice post RSV challenge. In contrast to the quick and massive body weight loss in FI-RSV immunized mice, body weight changes in P-KFD1 or P+CTB immunized mice were not significantly different from those in saline group after RSV challenge (**Figure S2A**). We further evaluated the respiratory function of mice by testing their airway hyper-responsiveness (AHR). As shown in **Figure 2A**, P-KFD1 immunized mice had normal inhalation resistance (Ri), exhalation resistance (Re) and lung dynamic compliance (Cdyn) after challenge, which were similar to those measured parameters in saline immunized mice and uninfected naïve mice. The FI-RSV immunized mice showed dramatic increases of both Ri and Re, and significant decrease of Cdyn by excitation with increasing concentrations of methacholine (MCH). The P+CTB immunized mice showed lower values of Ri and Re than the FI-RSV group, but higher values than those of either the P-KFD1 or saline groups. These results indicated that P-KFD1 intranasally immunized mice retained normal respiratory function after RSV challenge, in contrast to FI-RSV immunized mice which showed severe airway obstruction as reported in many studies (Knudson et al., 2015). In parallel with the respiratory function, we observed lung histopathology changes by H&E and PAS staining at 8 days post RSV infection (dpi) in naïve or immunized mice. As shown in **Figure 2B**, consistent with typical VED, overt inflammation and mucus production (black arrows) were observed in lungs of FI-RSV immunized mice. In contrast, neither inflammatory cell infiltration nor mucus secretion was observed in lungs of P-KFD1 immunized mice (**Figure 2B**). It should be noted that more inflammatory cell infiltration and mucus production were observed in the P+CTB immunized mice compared to the P-KFD1 group, but less compared to the FI-RSV group. In detail, the highest pathological scores of immunocyte aggregation around both bronchioles and pulmonary vessels, and the highest score of interstitial pneumonia were observed in FI-RSV immunized mice, while significant lower scores were observed in the P-KFD1 or saline immunized mice (**Figure 2C**). The P+CTB immunized mice showed lower scores of inflammations than the FI-RSV immunized mice, but higher scores than the P-KFD1 immunized mice. In addition, a significant lower score of mucus production was observed in the P-KFD1 immunized mice compared to that in either FI-RSV or P+CTB immunized mice (**Figure 2D**). Furthermore, analysis of differential infiltrated immunocytes in lungs by flow cytometry (FCM) showed an increase of eosinophils and T cells in the FI-RSV immunized mice, and an increase of T cells in the P+CTB immunized mice, compared to those of P-KFD1 or saline immunized mice upon RSV challenge (**Figure 2E**). We also analyzed the Th bias in lungs of the immunized mice at 8 days after challenge by *ex-vivo* PMA and ionomycin stimulation (**Figure 2F**). Compared to saline immunized mice, percentage of IL-4 secreting cells in CD4^+^ T cells increased significantly in FI-RSV immunized mice as many studies previously reported (Knudson et al., 2015), while no increase was observed in either the P-KFD1 or P+CTB immunized mice. Moreover, there was no difference in frequency of CD25^+^ Foxp3^+^ Treg (**Figure S2B**) or IFN-γ^+^ CD4^+^ T cells among all groups of mice. It should be noted that a significant elevated percentage of IL-17A^+^ CD4^+^ T cells was detected in P+CTB immunized mice. In summary, in contrast to FI-RSV immunization, i.n. immunization with P-KFD1 rather than P+CTB completely avoided occurrence of VED. Taken the anti-RSV efficacy in account (**Figure 1D**), P-KFD1 might be a potential safe and effective RSV vaccine candidate.

**Figure 2.**
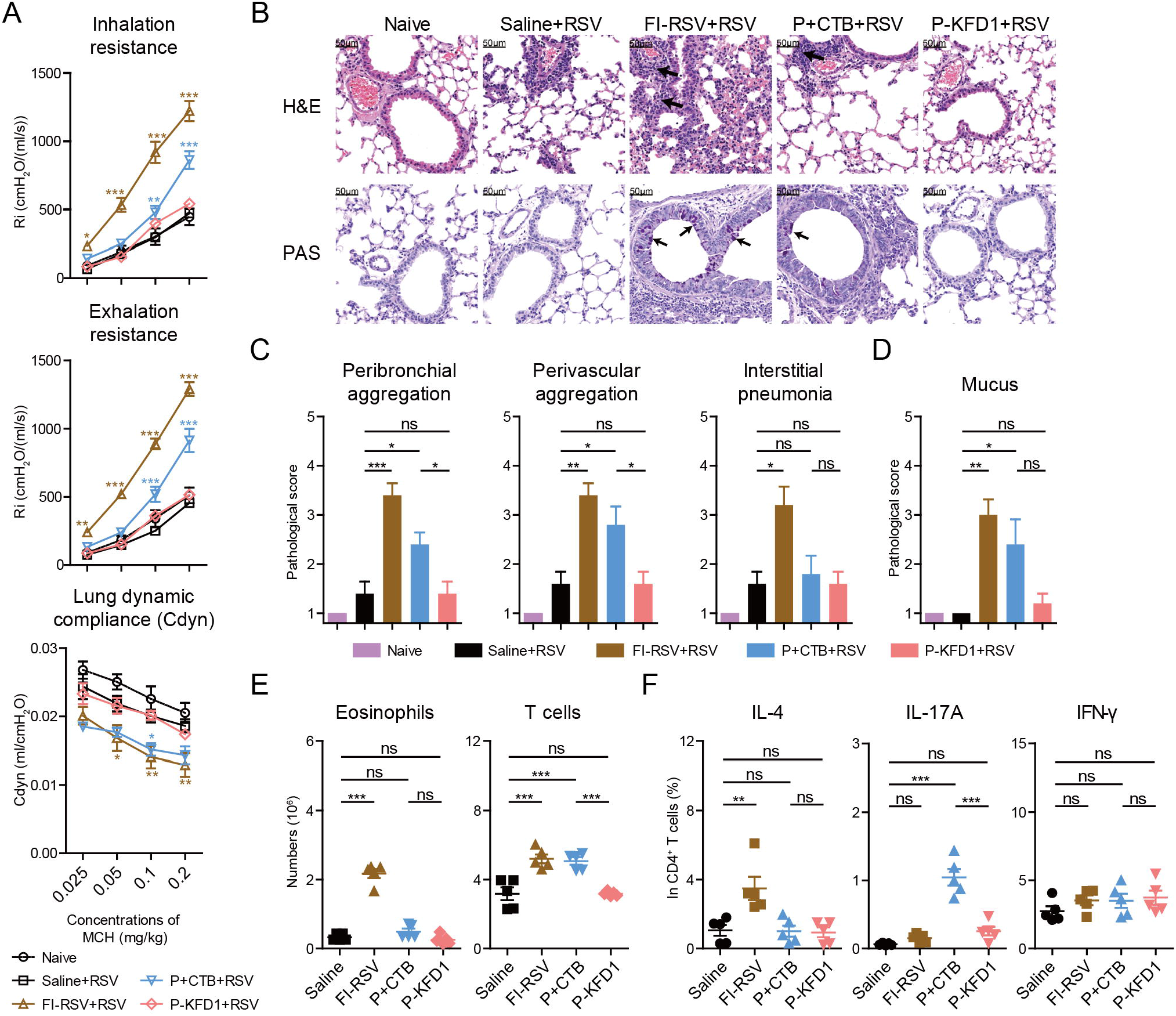
Intranasal immunization with P-KFD1 protects mice against lung disease caused by RSV infection. Groups of BALB/c mice were immunized with saline, FI-RSV, P+CTB or P-KFD1, respectively, prior to RSV challenge, and sacrificed on 8 dpi. (**A**) Inspiratory resistance, expiratory resistance and lung dynamic compliance of mice were monitored, n = 5 mice per group. (**B**) Hematoxylin and eosin (H&E) and periodic acid-Schiff (PAS) staining were performed on lung sections. (**C**) Pathological scores of immunocytes aggregation around bronchioles, pulmonary vessels and scores of interstitial pneumonias were evaluated according to the H&E-stained sections, n = 5 mice per group. (**D**) Scores of mucus production were evaluated according to the PAS-stained sections, n = 5 mice per group. (**E**) Numbers of eosinophils and T lymphocytes in lungs of mice were assessed by flow cytometry (FCM), n = 5 mice per group. (**F**) Percentages of IL-4^+^, IL-17A^+^ or IFN-γ^+^ cells in CD4^+^ T cells with PMA and ionomycin stimulation in lungs of mice were determined by FCM, n = 5 mice per group. Data are represented as mean ± SEM of two independent experiments. In **Figure 2A**, groups were compared using regular two-way ANOVA with saline group. In **Figures 2C to 2F**, groups were compared using one-way ANOVA. * p < 0.05, ** p < 0.01, *** p < 0.001, ns means non-significant. See also **Figure S2**.

### CD4^+^ T cells but not antibodies provide protection against RSV infection

The encouraging protective effects conferred by P-KFD1 i.n. immunization urged us to further investigate the relationship between protective efficacy and immunization strategy. We found that inoculation with 20 μg of P-KFD1 provided better protection against infection than with a dosage of 5 μg or 1.25 μg (**Figure S3A**), and boosting twice provided better protection against infection than once (**Figure S3B**). Hence, the protective efficacy of P-KFD1 i.n. immunization is dependent on both dosage of immunogen and times of immunization. These results suggested that the degree of immune responses induced by P-KFD1 i.n. immunization is correlated with protective efficacy. Then we tried to elucidate, whether humoral immune response, or cellular immune response, or both, mediated protection in P-KFD1 immunized mice.

Based on previous studies on the intracellular neutralization activities of anti-viral IgA *in vitro* and *in vivo* through pIgR mediated transcytosis (Burns et al., 1996; Yang et al., 2020; Zhou et al., 2019; Zhou et al., 2011), P-KFD1 i.n. immunization was performed in wild type mice, as well as pIgR knockout mice (pIgR^−/−^), to test if P-specific secretory IgA plays a role or not. In line with reported data (Asahi et al., 2002; Shimada et al., 1999), our results showed IgA antibodies accumulated in serum, but could not actively be transported to the mucosal surface by the lack of pIgR (**Figure S3C**). In parallel, P-KFD1 i.n. immunization induced lower level of P-specific serum IgG, but higher level of serum IgA and undetectable P-specific saliva IgA in pIgR^−/−^ mice compared to those in wild type mice (**Figure 3A**). Despite the striking difference in P-specific secreting antibody response, the protection provided by P-KFD1 i.n. immunization against RSV infection in both noses and lungs of pIgR^−/−^ mice was as similar as that of wild type mice (**Figure 3B**). This result suggested that P-specific mucosal IgA antibodies did not mediate the protection against RSV by intraepithelial neutralization related to pIgR. We then performed anti-serum transfer experiment and found no anti-viral efficacy could be conferred to naïve recipient mice by transfer of serum from the P -KFD1 immunized mice (**Figure 3C**). It suggested that P-specific antibodies in serum were not the main protective factors. Collectively, P-specific humoral immune response induced by P-KFD1 i.n. immunization does not appear to be the main mediator for protection against RSV infection.

**Figure 3.**
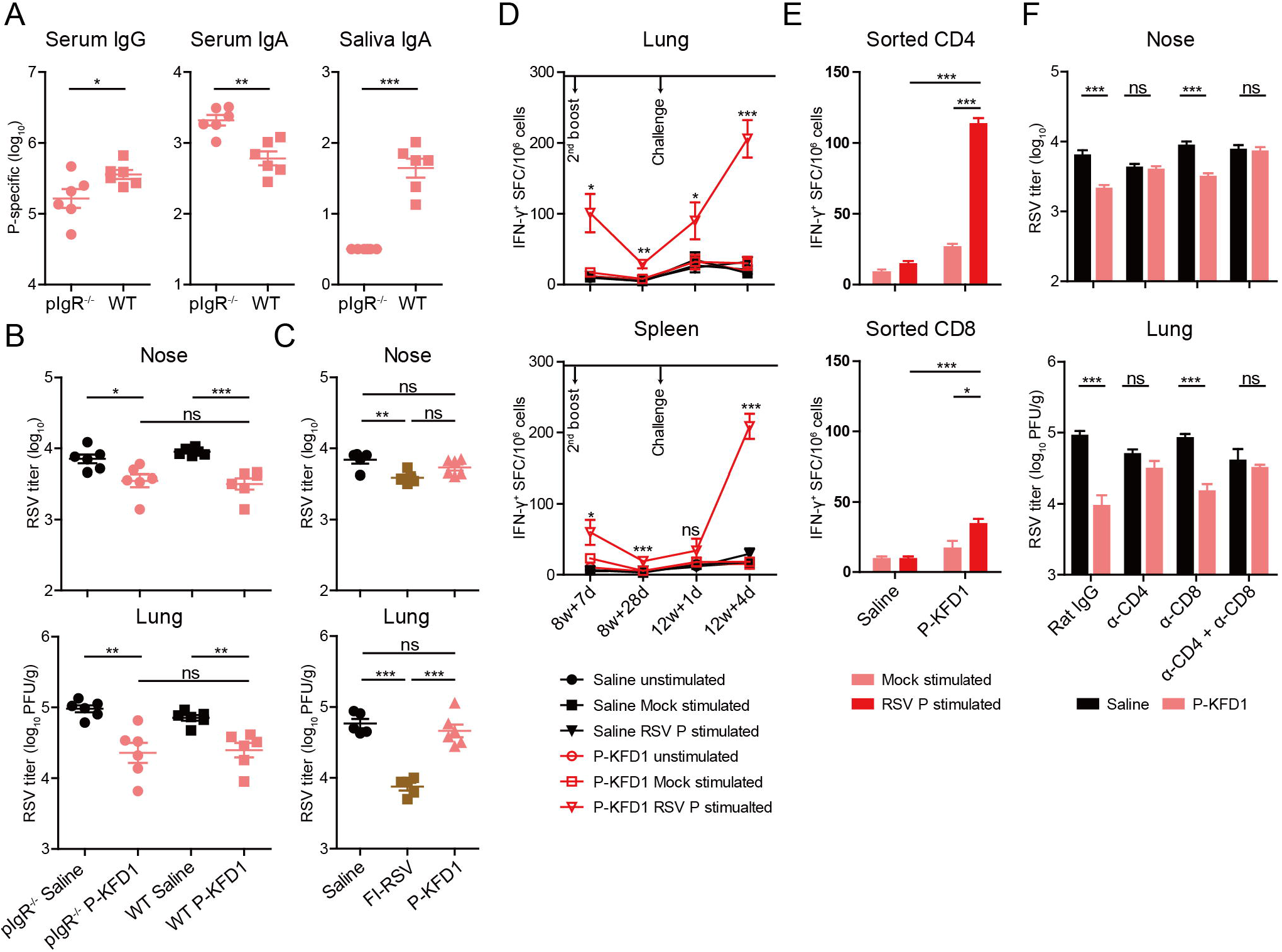
CD4^+^ T cells but not antibodies provide protection against RSV infection. (**A**) P-specific antibody responses in P-KFD1 immunized pIgR^−/−^ mice were measured at 14 days post last immunization, n = 6 mice per group. (**B**) Viral loads in saline or P-KFD1 immunized pIgR^−/−^ and wild type mice were measured on 4 dpi, n = 6 mice per group. (**C**) RSV titers in recipient mice that received immune sera from saline, FI-RSV or P-KFD1 immunized mice before RSV challenge, were monitored on 4 dpi, n = 5~6 mice per group. (**D**) P-specific IFN-γ^+^ T cell responses were determined by ELISpot at indicated time points, n = 5 mice per group. (**E**) P-specific IFN-γ^+^ responses of purified splenic CD4^+^ T cells and CD8^+^ T cells were detected by ELISpot on 7 days post last immunization, n = 4 replicates per group. (**F**) RSV titers in saline or P-KFD1 intranasally immunized mice that were treated with depletion antibodies against CD4^+^ and/or CD8^+^ T cells were detected on 4 dpi, n = 5~6 mice per group. Data are represented as mean ± SEM of two independent experiments. In **Figures 3A and 3B**, two groups were compared using un-paired t test. In **Figure 3C**, groups were compared using one-way ANOVA. In **Figures 3D to 3F**, groups were compared using regular two-way ANOVA. * p < 0.05, ** p < 0.01, *** p < 0.001, ns means non-significant. See also **Figure S3**.

Thus, we speculated that the T cell response induced by P-KFD1 i.n. immunization do mediate protection. As showed in **Figures 1B and S3D**, P-KFD1 i.n. immunization induced P-specific IFN-γ, IL-17A but not IL-4 secreting T cell responses. P-specific IFN-γ secreting T cells could be detected upon stimulation with P in both lungs and spleens at day 7 post second boost immunization (8w+7d), decreasing to nearly undetectable levels thereafter (8w+28d), but starting to rebound after challenge at 1 dpi (12w+1d) and increasing to a still higher level at 4 dpi (12w+4d) (**Figure 3D**). Accordingly, after challenge, the virus replicated only from 1×10 ^3^ at 1 dpi to 1.5×10 ^3^ at 4 dpi in lungs of P-KFD1 immunized mice, but replicated dramatically from 0.5×10 ^3^ at 1 dpi to 1×10 ^5^ at 4 dpi in lungs of saline immunized mice (**Figure S3E**). This delayed anti-virus effect implied that T cells might take part in the protection conferred by P-KFD1 i.n. immunization. With regard to IFN-γ production by purified CD4^+^ or CD8^+^ T cells, P-KFD1 i.n. immunization induced much higher fraction of P-specific IFN-γ producing CD4^+^ T cells than the fraction of P-specific IFN-γ producing CD8^+^ T cells (**Figure 3E**). This indicated that the P-specific CD4^+^ T cells elicited by P-KFD1 i.n. immunization should be the main cellular source of IFN-γ.

To confirm the role of T cells played in the protection against infection, CD4^+^ and/or CD8^+^ T cells were depleted *in vivo* by using anti-CD4-specific and/or anti-CD8-specific antibodies (**Figure S3F)**. When T cells were depleted simultaneously with anti-CD4 and anti-CD8 antibody together (α-CD4 + α-CD8), the anti-viral activity in P-KFD1 immunized mice was abrogated in both noses and lungs (**Figure 3F**). When only CD4^+^ T cells were depleted, the anti-viral efficacy in P-KFD1 immunized mice declined significantly to the same level as that in saline immunized mice. However, if only CD8^+^ T cells were depleted, anti-viral efficacy remained intact in both noses and lungs (**Figure 3F**). Hence, the protection against RSV conferred by i.n. immunization with P-KFD1 was mainly CD4^+^ T cell dependent.

### Both local and peripheral CD4^+^ T cells play roles in protection against RSV infection

Considering the importance of local immunity for protection against RSV infection in the upper respiratory tract and lung, we further investigated whether the protective CD4^+^ T cells resided in respiratory tract or migrated from lymphoid organ. To make this determination, a sphingosine-1-phosphate receptor 1 agonist, FTY720, was adopted for blocking the egress of T lymphocytes from lymphoid organs of experimental mice (Brinkmann et al., 2002; Mandala et al., 2002). As shown in **Figure 4A**, FTY720 treatment did not affect the number of P-specific IFN-γ producing T cells in spleens of P-KFD1 immunized mice, but rather decreased those in lungs significantly from about 200 to 70 SFC/10^6^ lymphocytes. This result indicated that in lung post infection, more than half of the P-specific IFN-γ producing T cells migrated from lymphoid organs, and could be blocked by FTY720 treatment. Consistent with the decreased migration of T cells, the anti-viral efficacy of P-KFD1 immunized mice was mitigated by the FTY720 treatment, while no distinguishable difference was observed in saline immunized mice, irrespective of FTY720 treatment (**Figure 4B**). This suggested circulating T cells took part in protection afforded by P-KFD1 i.n. immunization. Meanwhile, FTY720 treatment just partially abrogated the anti-viral efficacy of the P-KFD1 immunized mice, in which viral titers in both noses and lungs were still significantly lower than those of the saline immunized mice, also treated with FTY720 (**Figure 4B**), further demonstrating that resident T cells in the lungs also played a protective role in P-KFD1 i.n. immunization. On the other hand, compared with i.n. immunization, intraperitoneal (i.p.) immunization with P-KFD1 induced comparable levels of P-specific IFN-γ^+^ T cell response and production of IFN-γ in the spleen, but lower levels in lung (**Figure S4**). In addition, i.p. immunization with P-KFD1 resulted in significantly reduced viral load in lung, but not in nose (**Figure 4C**). Taken together, both the migrated T cells from lymphoid organ and the resident T cells in the respiratory tract contributed to local *in situ* anti-viral immunity in P-KFD1 immunized mice.

**Figure 4.**
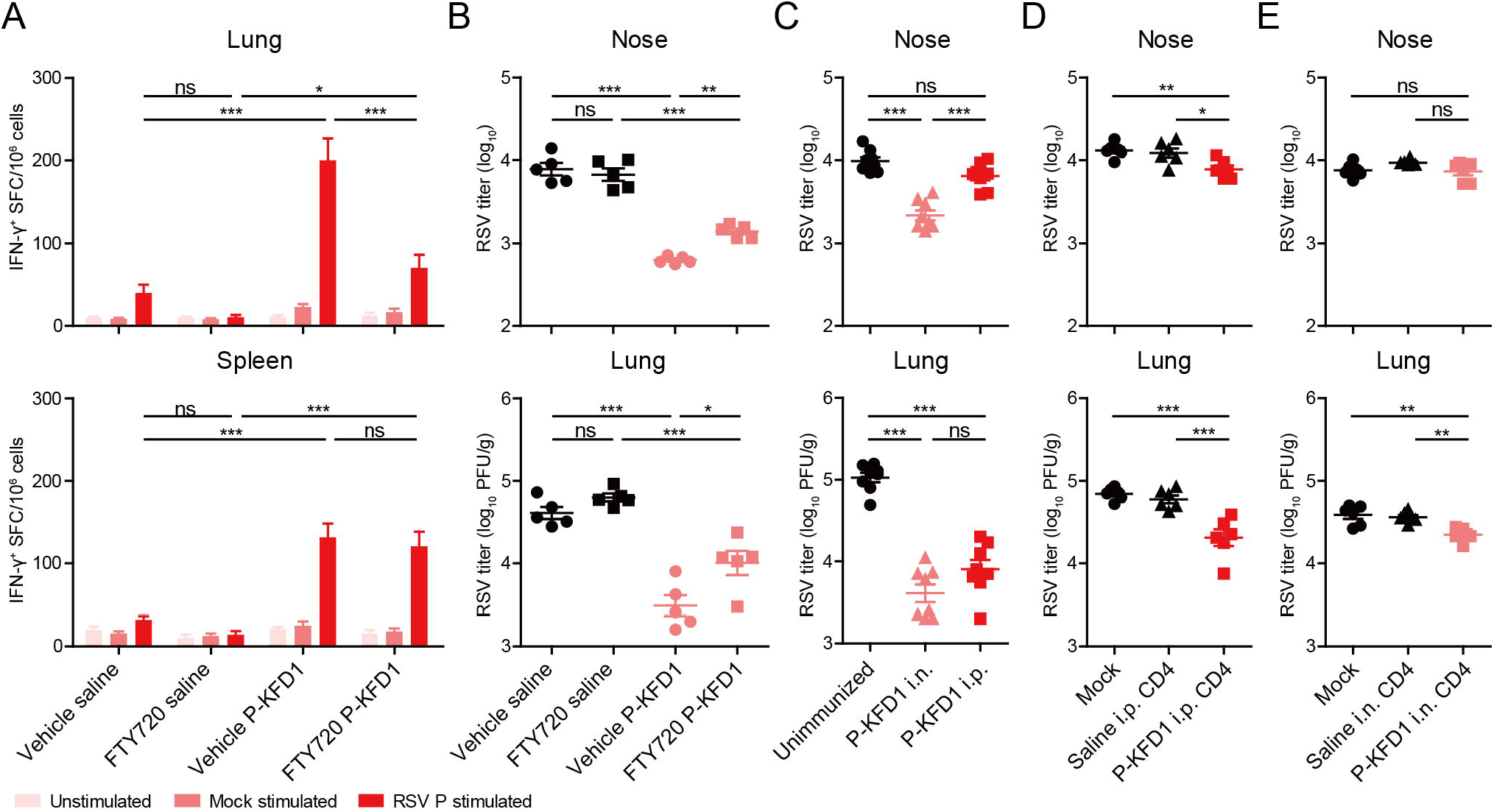
Both local and peripheral CD4^+^ T cells play roles in protection against RSV infection. (**A and B**) Saline or P-KFD1 intranasally immunized mice were intraperitoneally injected with vehicle or FTY720 daily on 7 successive days. (**A**) P-specific IFN-γ^+^ T cell response in lungs and spleens (n = 10 mice per group) and (**B**) RSV titers in noses and lungs of mice (n = 5 mice per group) were detected at 4 dpi. (**C**) Mice were intranasally or intraperitoneally immunized with P-KFD1 at four-week intervals three times prior to RSV challenge. RSV titers in noses and lungs were determined at 4 dpi, n = 8 mice per group. (**D and E**) Splenic CD4^+^ T cells were sorted by microbeads from saline or P-KFD1 immunized mice and were intravascularly injected into naïve mice, respectively, followed by RSV challenge. Viral loads in noses and lungs of recipients that received CD4^+^ T cells from intraperitoneally immunized donors (**D**) or from intranasally immunized donors (**E**) were detected at 4 dpi, n = 6 mice per group. Data are represented as mean ± SEM of two pooled experiments for **Figures 4A and 4C**. Data are represented as mean ± SEM of two independent experiments for **Figures 4B, 4D and 4E**. In **Figure 4A**, groups were compared using regular two-way ANOVA. In **Figures 4B to 4E**, groups were compared using one-way ANOVA. * p < 0.05, ** p < 0.01, *** p < 0.001, ns means non-significant. See also **Figure S4**.

To further verify that it was CD4^+^ T cells evoked by P-KFD1 immunization that directly contributed to protection against viral infection, we performed adoptive transfer experiments. Briefly, CD4^+^ T cells from the spleens of intraperitoneally or intranasally immunized mice were sorted by antibody-conjugated microbeads and then transferred into naïve mice prior to RSV challenge. Mice that received CD4 ^+^ T cells from P-KFD1 intraperitoneally immunized donors had lower viral loads in both noses and lungs post challenge (**Figure 4D**). Mice that received CD4^+^ T cells from P-KFD1 intranasally immunized mice also exhibited lower viral load in lungs, but only marginally lower viral titer in the noses (**Figure 4E**). Collectively, these data suggested that the P-specific CD4^+^ T cells induced by P-KFD1 i.n. immunization, either resided in, or migrated to the respiratory tract, played key protective roles against RSV infection.

### Sc-RNA seq reveals a cluster of specific Th17 cells corresponding to the P-KFD1 i.n. immunization

Next, we tried to directly investigate the CD4^+^ T cell responses elicited by P-KFD1 i.n. immunization during the acute phase of RSV infection by sc-RNA seq (single cell RNA sequencing). As cell surface integrin molecules CD11a and CD49d can be used to distinguish antigen-experienced CD4^+^ T cells and naïve CD4 ^+^ T cells (Knudson et al., 2014; McDermott and Varga, 2011), we sorted the antigen-experienced CD11a^high^ CD4^+^ T cell subset and antigen-unexperienced CD11a^low^ CD49d^−^ CD4^+^ T cell subset from lungs of saline or P-KFD1 intranasally immunized mice on 4 dpi by FACS (**Figures 5A and S5A**). As expected, P-specific IFN-γ^+^ CD4^+^ and IL-17A^+^ CD4^+^ T cell responses could only be detected in the CD11a^high^ CD4^+^ T cell subsets of P-KFD1 immunized mice (**Figure S5B**). To focus on these responsive CD11a^high^ cells in the CD4^+^ T cells, we mixed the sorted CD11a^high^ subset with hashtag antibody labeled sorted CD11a^low^ CD49d^−^ cell subset at a ratio of approximately 5:3, before performing sc-RNA seq of the transcriptome profile and TCR repertoire of CD4^+^ T cells (**Figure 5B**). Unsupervised hierarchical clustering and t-distributed stochastic neighbor embedding (t-SNE) dimensional reduction analysis based on the transcriptomes of pooled samples identified fifteen CD4^+^ T cell clusters (**Figures 5C and S5C**). The cells in clusters 1, 3 and 7 were naïve -like CD4^+^ T cells, since they expressed transcripts of *Ccr7*, *Lef1* and *Igfbp4* (**Figures 5D and S5E**). Cells in cluster 0 highly expressed transcripts of *Itgb1*, *Cd40lg*, *Il2*, and *Ifng*, which resembled effector memory or tissue resident memory CD4^+^ T cells (Tem/Trm) (**Figure 5D**). Whereas, cluster 4 cells highly expressed transcripts of *Il2ra, Foxp3* and *Ikzf2* which seemed like Treg cells. Notably, cluster 6 cells were enriched in *Il17a*, *Il17f*, *Ccr6* and *Rorc*, indicative of Th17 subset (**Figure 5D**). Other clusters were also designated based on the signature genes expression (**Figure 5D**). Moreover, sc-RNA seq revealed a significantly higher proportion of Th17 cells in P-KFD1 induced CD4^+^ T cells compared to saline group (**Figure 5C**).

**Figure 5.**
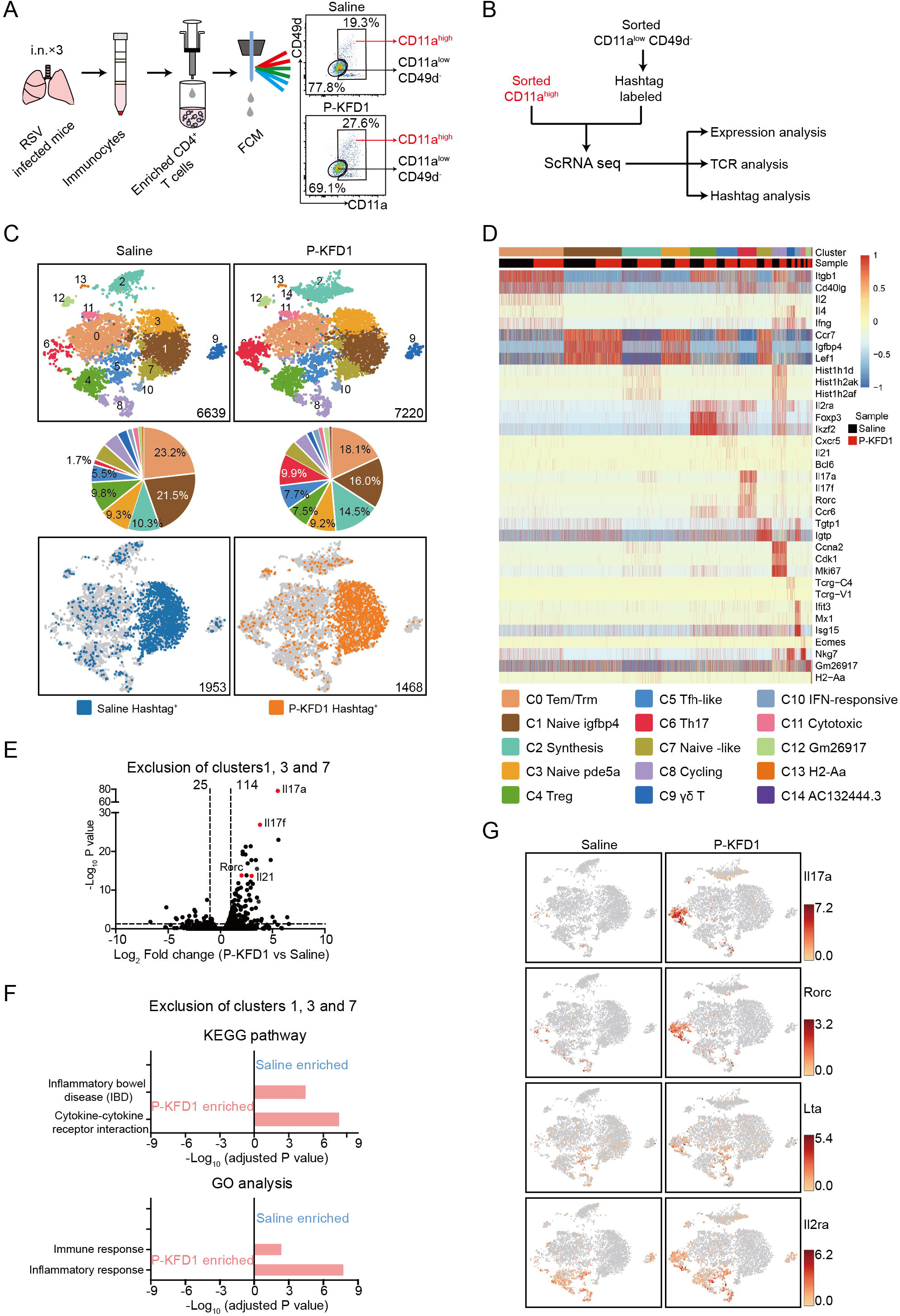
Sc-RNA seq of CD4^+^ T cells reveals the elevation of activation- and cytotoxicity-associated genes. (**A**) Schematic diagram of isolating CD11a^high^ CD4^+^ and CD11a^low^ CD49d^−^ CD4^+^ T cell subsets in saline or P-KFD1 intranasally immunized mice on 4 dpi. (**B**) Overview of sc-RNA seq of pulmonary CD4^+^ T cells. FACS sorted CD11a^low^ CD49d^−^ CD4^+^ T cells were labeled by hashtag antibody and mixed with CD11a^high^ CD4^+^ T cells at a ratio of approximately 3:5 before sc-RNA seq of mRNA and TCR. (**C**) Clustering results and hashtag analysis of sc-RNA seq. Each point represents one single barcode labeled cell and each cluster was displayed in the t-SNE plots established by total mRNA. Pie charts exhibited percentages of detected barcodes in each cluster. (**D**) Heatmap of clustering identification associated signature differential expression genes (DEGs) in each cluster. Each vertical line represents a single cell. (**E**) Volcano plot of DEGs in cells exclusion of clusters 1, 3 and 7 between P-KFD1 group versus saline group. (**F**) Enriched KEGG pathways and GO terms for DEGs in non-naive cells excluded clusters 1, 3 and 7 between P-KFD1 group versus saline group. (**G**) T-SNE plots for gene expression of DEGs *Il17a*, *Rorc*, *Lta* and *Il2ra*. See also **Figure S5**.

As expected, the hashtag labeled antigen-unexperienced CD11a^low^ CD49d^−^ CD4^+^ T cells were mainly located in the naïve -like clusters 1, 3 and 7 cells (**Figures 5C and S5D**), and showed genes expression pattern similar as naïve CD4 ^+^ T cells (**Figure S5E**). Moreover, similar as that in the naïve -like cells, differential expression genes (DEGs, log_2_|FC| > 1, P<0.05) were rarely found in hashtag^+^ cells between P-KFD1 group versus saline group (**Figure S5F**). In contrary, large amount of DEGs (139 genes) could be identified in the non-naïve cells that excluded clusters 1, 3 and 7 in P-KFD1 group compared to saline group (**Figure 5E**). David analysis revealed that the upregulated DEGs in P-KFD1 group were enriched in two KEGG pathways “cytokine-cytokine receptor interaction” and “inflammatory bowel disease”, and in two Gene Ontology (GO) biology processes “inflammatory response” and “immune response” in P-KFD1 group (**Figure 5F**). The representative DEGs included Th17 marker genes *Il17a*, *Il17f, Rorc*, *Ccr6*, activation related marker genes *Il2ra, Cxcr6*, and cytotoxicity-associated genes *Lta* etc. (**Figures 5G and S5G**).

Then we aimed at the set of T cells with the same TCRs, which were the sequencing identified clonal expanded CD4^+^ T cells. TCR sequencing totally identified 5695 and 5769 barcodes in saline group and P-KFD1 group, respectively. It is obviously that the frequency and the number of T cells with repeatedly used TCRs in P-KFD1 group (625) were much higher than that in saline group (330) (**Figures 6A, S6A and S6B**). Moreover, the clone sizes of CD4^+^ T cells in P-KFD1 group were much larger than that in saline group (**Figure 6B**). In addition, in P-KFD1 group, a large portion of the CD4^+^ T cells with repeatedly used TCRs were located in cluster 6 Th17 cells. These suggested epitope-specific CD4^+^ T cell responses especially the Th17 cells were elicited by P-KFD1 i.n. immunization. Consistently, between saline group and P-KFD1 group, TCR usage in the non-naïve CD4^+^ T cells that excluded clusters 1,3,7 or cells with repeatedly used TCRs were significantly different (**Figures S6C and S6D**). In CD4^+^ T cells with repeatedly used TCRs, large amount of DEGs existed between P-KFD1 group versus saline group (**Figure 6C**). These upregulated genes in P-KFD1 group also included Th17 related transcripts of genes (*Ccr6*, *Il17a*, *Il17f* and *Il23r*), activation related genes such as *Il2ra* and *Cxcr6*, as well as some immune response-associated genes such as *Ccr1*, *Csf1*, *Lta*, *Il1r1* and *Tnfsf11* (**Figures 6D and S6E**). The results above all suggested that P-KFD1 i.n. immunization induced expansion and activation of Th17 cells on 4 dpi at transcriptional level.

**Figure 6.**
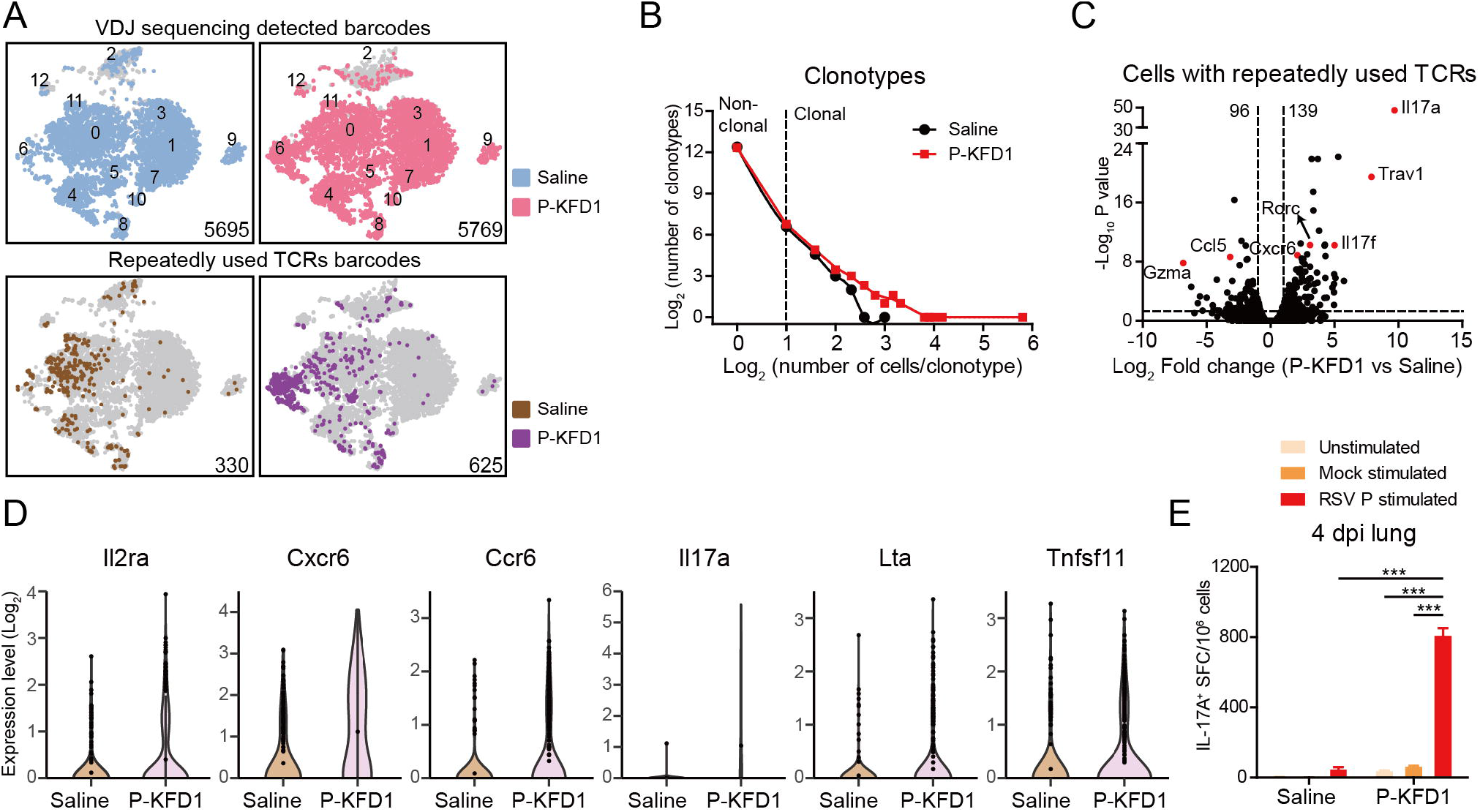
Clonal expansion of Th17 cells and P-specific IL-17A^+^ T cell response in P-KFD1 intranasally immunized mice on 4 dpi. **(A-D)** TCR sequencing analysis. (**A**) T-SNE plots for TCR sequencing detected barcodes (upper panel) and repeatedly used TCRs barcodes (lower panel). (**B**) The association between the number of CD4^+^ T cell clonotypes and the number of cells per clonotype in saline and P-KFD1 groups. (**C**) Volcano plot of DEGs in cells with repeatedly used TCRs between P-KFD1 group versus saline group. (**D**) Violin plots of some significant DEGs in cells with repeatedly used TCRs. (**E**) ELISpot assay of P-specific IL-17A^+^ T cell response of pulmonary immunocytes on 4 dpi. n = 6 mice per group. Data are represented as mean ± SEM of at least two independent experiments for **Figure 6E**, and groups were compared using regular two-way ANOVA. *** p < 0.001. See also **Figure S6.**

ELISpot assay of pulmonary immunocytes showed that much higher levels of P-specific IL-17A^+^ T cell response were elicited in P-KFD1 intranasally immunized mice on 4 days after RSV challenge (**Figure 6E**). Generally, P-KFD1 i.n. immunization boosted P-specific Th17 immune responses.

### P-specific IL-17A cooperates with IFN-γ in resisting RSV infection

Then, transfer model was adopted to further confirm the characteristics of CD4^+^ T cell responses induced by P-KFD1 i.n. immunization. Briefly, splenocytes prepared from P-KFD1, KFD1 or saline intranasally immunized mice were labeled with carboxyfluorescein succinimidyl ester (CFSE) and intravascularly injected into naïve mice at 12 hours prior to RSV challenge respectively. Lymphocytes in lungs and spleens of the recipient mice were isolated at 4 days post challenge and analyzed by FCM (**Figure 7A**). The CFSE-labeled cells derived from donor mice could be detected and differentiated as un-proliferated CFSE^high^ cells and proliferated CFSE^low^ cells (**Figure 7B**). As depicted in **Figure 7C**, in the CFSE^+^ cells, percentages of CFSE^low^ CD4^+^ T cells from P-KFD1 immunized mice were significantly higher than those from KFD1 or saline immunized donors, in both lungs and spleens of the recipient mice, but CFSE^low^ CD8^+^ T cells were not. This result indicated that P-specific CD4^+^ T cells induced by P-KFD1 i.n. immunization could expand upon RSV challenge.

**Figure 7.**
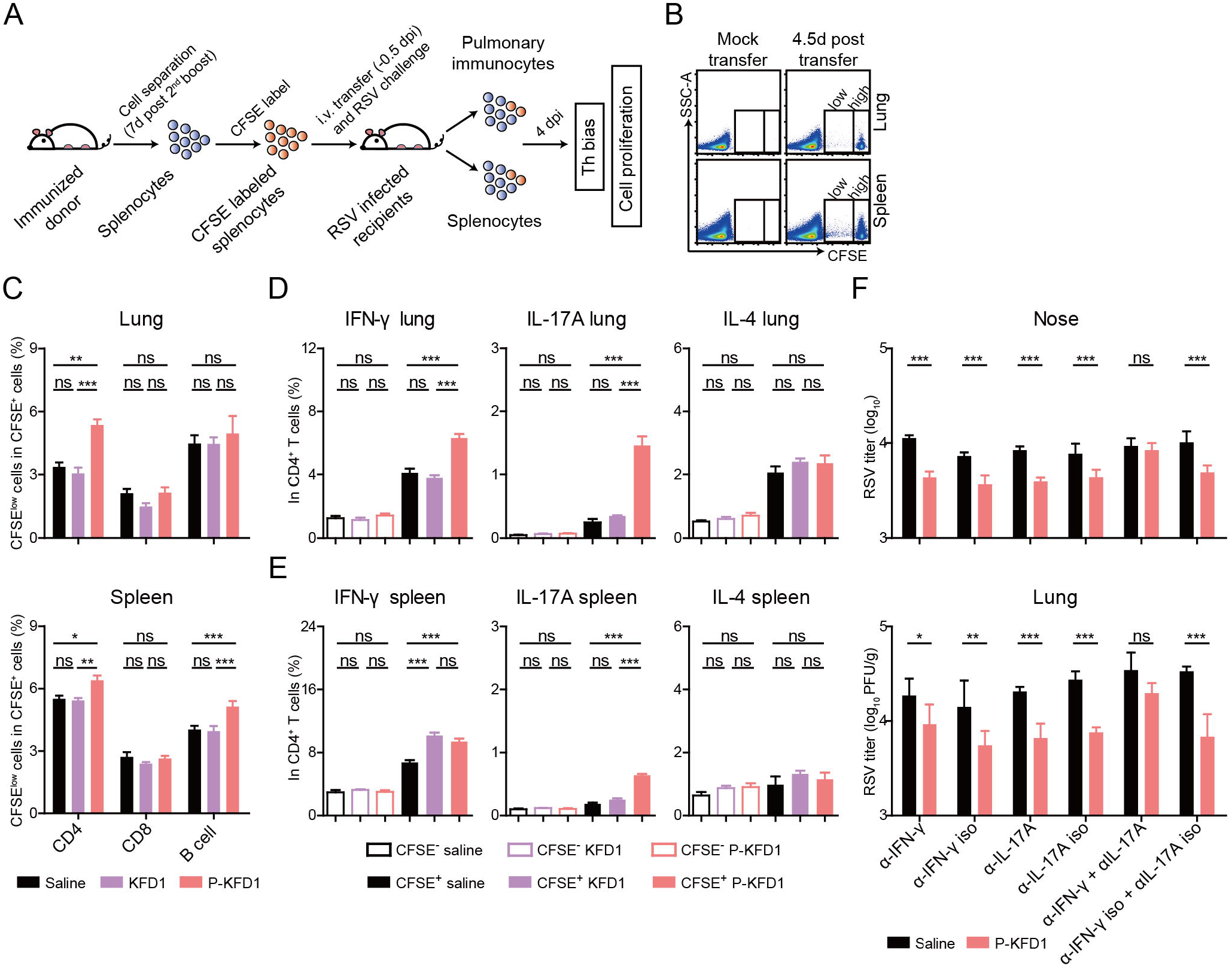
P-specific Th1 and Th17 immune responses induced by P-KFD1 i.n. immunization contribute to protection. (**A to E**) Adoptive transfer experiment. (**A**) Flow diagram of adoptive transfer experiment. (**B**) Representative plots of immunocytes in lungs and spleens of recipient mice with or without CFSE-labeled cells. (**C**) Percentages of proliferated CD4^+^, CD8^+^ T cells and B cells from immunized donors in lungs and spleens of recipients were evaluated at 4 dpi, n = 5 mice per group. (**D and E**) With PMA and ionomycin stimulation, percentages of IFN-γ^+^, IL-17A^+^ or IL-4^+^ cells in CFSE^−^ CD4^+^ or CFSE^+^ CD4^+^ T cells in lungs (**D**) and spleens (**E**) of recipients were assessed at 4 dpi, n = 4~5 mice per group. (**F**) Saline or P-KFD1 intranasally immunized mice were treated with antibodies to block IFN-γ, IL-17A or both IFN-γ and IL-17A during RSV infection, respectively, viral loads were assessed on 4 dpi, n = 4~5 mice per group. Data are represented as mean ± SEM of at least two independent experiments. In **Figures 7C and 7F**, groups were compared using regular two-way ANOVA. In **Figures 7D and 7E**, groups were compared using one-way ANOVA. * p < 0.05, ** p < 0.01, *** p < 0.001, ns means non-significant. See also **Figure S7**.

In order to detect potential atopy of Th responses, lymphocytes of the recipient mice were stimulated by PMA and ionomycin *ex vivo* and analyzed. The percentages of IFN-γ^+^ cells, IL-17A^+^ cells or IL-4^+^ cells in CFSE^−^ CD4^+^ T cells did not differ among the three different recipient groups in lungs and spleens, respectively, suggesting that adoptive transfer did not change Th bias of recipient mice after RSV challenge (**Figures 7D and 7E, hollow column**). In CFSE^+^ CD4^+^ T cells, both the percentages of IFN-γ^+^ cells and IL-17A^+^ cells from P-KFD1 immunized donors were significantly higher than those from saline immunized donors, in lungs and spleens of recipients respectively. Whereas, no differences were found in percentages of CFSE^+^ IL-4^+^ CD4^+^ T cells among all groups (**Figures 7D and 7E, solid column**). These results indicated P-KFD1 i.n. immunization induced both Th1 and Th17 responses at 4 dpi. We also found that the percentages of CFSE^+^ IFN-γ^+^ CD8^+^ T cells from P-KFD1 intranasally immunized donors were slightly higher than those from saline immunized donors (**Figure S7A**). However, the percentages of CFSE^+^ IL-17A^+^ CD8^+^ T cells were all at low levels and showed no difference among the recipients (**Figure S7A**).

Furthermore, the percentages of CFSE^+^ IFN-γ^+^ CD4^+^ T cells, CFSE^+^ IL-17A^+^ CD4^+^ T cells (**Figure 7D**) and CFSE^+^ IFN-γ^+^ CD8^+^ T cells (**Figure S7A**) from P-KFD1 immunized donors were higher than those from KFD1 immunized mice.

These results implied that P-KFD1 i.n. immunization induced P-specific Th1 and Th17 responses, as well as IFN-γ^+^ CD8^+^ T cell response, especially in RSV infected tissue. When the same CFSE labeling and adoptive transfer experiments were performed for the FI-RSV and P+CTB immunized mice, a dramatically higher level of CD4^+^ T cell proliferation at 4 dpi could be observed compared to that of P-KFD1 immunized mice (**Figure S7B**). Besides, a robust Th2-biased response in the FI-RSV immunized mice and a robust Th17-biased immune response in the P+CTB immunized mice could be observed in both lungs and spleens, respectively (**Figure S7C**), which were also consistent with the results in **Figure 2F**.

To determine whether the Th1 and/or Th17 responses mediated anti-viral efficacy in P-KFD1 intranasally immunized mice, antibodies against IFN-γ (α-IFN-γ) and/or IL-17A (α-IL-17A) were injected into the P-KFD1 immunized mice during RSV infection, respectively. As shown in **Figure 7F**, simultaneous administration of both anti-IFN-γ and anti-IL-17A antibodies could abolish anti-viral efficacy in both noses and lungs of P-KFD1 intranasally immunized mice. In contrast, administration of only anti-IFN-γ antibody or anti-IL-17A antibody could not abrogate anti-viral efficacy; instead, significantly lower viral loads were retained in both noses and lungs of P-KFD1 intranasally immunized mice compared to those of saline immunized mice, respectively. These data suggested that T cell-derived IFN-γ or IL-17A alone could mediate the inhibition of RSV replication *in vivo*. Taken together, P-KFD1 i.n. immunization induced P-specific IFN-γ^+^ CD4^+^ and IL-17A^+^ CD4^+^ T cell responses without changing Th bias, and provided protection against RSV infection depended on IFN-γ or IL-17A.

## DISCUSSION

In past decades, many efforts and resources have been devoted to the development of a safe and efficacious RSV vaccine to prevent infection by this respiratory pathogen. However, the fact that 1) natural RSV infection does not elicit long-lasting immunity and 2) repeated infections occur throughout life (Glezen et al., 1986; Henderson et al., 1979) results in a dilemma for investigators designing a vaccination strategy by mimicking the host immune response, following natural infection with RSV. Furthermore, VED triggered by immunization with FI-RSV has presented yet another major obstacle against the development of an RSV vaccine (Kapikian et al., 1969; Kim et al., 1969).

Here, we reported how we used new strategies to develop a safe and effective mucosal RSV vaccine targeting an internal RSV antigen P that has been seldom investigated. And we examined the immune responses in relation to virological and pathological features of the fusion protein P-KFD1 intranasally immunized animals before and after RSV challenge by comparison with those immunized with P+CTB or FI-RSV. Our data showed that P-KFD1 i.n. immunization induces P-specific immune responses and inhibits RSV replication in both the upper and lower respiratory tract of mice. It is noteworthy that P-KFD1 i.n. immunization does not result in VED, whereas P+CTB i.n. immunization do cause some enhanced respiratory disease (ERD) which is different from the typical VED featured by the infiltration of eosinophils and highly Th2-biased immune response in FI-RSV immunized mice.

As part of our initial hypothesis, P-specific secreted IgA response in P-KFD1 immunized mice would play a role in prevention against viral infection. However, neither P-specific secreted IgA nor P-specific serum was found to be relevant to protection against RSV infection in our immunization regimen, though our experiments performed with pIgR^−/−^ mice, or by serum transfer, could not completely exclude the possible roles of antibodies or B cells through other unknown mechanisms.

We thus analyzed the kinetics of T cell response associated with P-KFD1 i.n. immunization and RSV challenge. We demonstrated that P-specific CD4^+^ T cells, either in lung or spleen, induced by P-KFD1 i.n. immunization played a central role in the protection against RSV infection. Our finding on the protective relevance of the RSV P-specific CD4^+^ T cells in mouse model is in line with the preexisting influenza-specific memory CD4^+^ T cells in blood, which were found to be correlated with disease protection against influenza challenge in humans (Wilkinson et al., 2012). Studies on influenza and coronavirus further showed the antigen-specific CD4^+^ T cells can resist infection by releasing IFN-γ (McKinstry et al., 2012; Zhao et al., 2016). A subcutaneous immunization with recombinant BCG strain expressing RSV N could also induce protective Th1-type immunity against RSV infection in mice (Bueno et al., 2008; Cautivo et al., 2010). These previous studies further support our finding on the protective role of P-specific Th1 immune response induced by P-KFD1 i.n. immunization.

The role of Th17 response remains controversial in different circumstances. A growing body of evidence indicates that Th17 cells play an important role in resisting some bacterial and fungal infections (Bacher et al., 2019; Li et al., 2018; Shao et al., 2019). An influenza virus vaccine candidate, termed CTA1-3M2e-DD, induced M2e-specifc memory CD4^+^ T cells that protected against infection also in an IL-17-dependent manner (Eliasson et al., 2018). However, the Th17 immune response is usually considered pathogenic by the tendency to exacerbate inflammatory response, which is closely related to several diseases, such as chronic obstructive pulmonary disease, cystic fibrosis and asthma (Iwanaga and Kolls, 2019). IL-17 produced during RSV infection has also been reported to be involved in increasing mucus secretion (Mukherjee et al., 2011). With the help of sc-RNA seq, we clearly identified the Th17 expansion in P-KFD1 intranasally immunized mice post infection. Moreover, besides Th17 representative genes (such as *Rorc*, *Ccr6*, *Il17a* and *Il17f*) and activation related genes (such as *Il2ra* and *Cxcr6)*, some cytotoxicity-associated genes such as *Lta*, and *Tnfsf11* were also significantly upregulated. Future investigation to determine whether these cytotoxicity-associated genes contribute to the protection against RSV infection is warranted. In our adoptive transfer experiments, both Th1 and Th17 immune responses were detected in the P-KFD1 and P+CTB groups, but the magnitude of Th17 immune response in the P+CTB group was much higher than that in the P-KFD1 group. A kind of aberrant Th17 response might explain why impaired respiratory function, inflammatory cell infiltration and mucus production occurred in the P+CTB immunized and RSV challenged mice. The underlying mechanism also needs further investigation.

Th1-related immune responses may also contribute to immunopathological damage during respiratory virus infection under some circumstances (Knudson et al., 2015; Wimalasundera et al., 1997). For any candidate RSV vaccine, safety is as important as effectiveness. In this study, P-KFD1 i.n. immunization induced moderate P-specific Th1 and Th17 immune responses without affecting Th bias, which might be involved in averting adverse effects. Although subunit vaccines are not the strongest vaccine format to induce T cell responses, i.n. immunization with P-KFD1 did induce T cell responses in a manner that struck a balance between safety and effectiveness in the typical RSV infection mouse model. Using other immunization strategies to elevate the magnitude of T cell responses might, however, change Th bias and induce ERD, such as the symptoms that appeared in P+CTB immunized mice, as discussed above.

Our animal model is a limiting factor in this study. Mice are semi-permissive for RSV replication. BALB/c mice infected with commonly used laboratory RSV strains, including the A2 strain we used, do not exhibit high viral loads or pulmonary mucus (Meng et al., 2014). Nevertheless, either viral load or pulmonary mucus production was measurable for comparatively evaluating both anti-viral efficacy and immunopathology in the mice subjected to different vaccinations in our study. Another caveat is that although the Th17 immune response in P-KFD1 intranasally immunized mice were characterized, the detail protective mechanism remains elusive. Further investigation on the mechanism and function of Th17 response during RSV infection is warranted to strike an optimal balance between anti-viral activity and immunopathology. Finally, we asked if P-specific T cell responses could be boosted to afford greater protection against RSV infection in human beings, even though researchers (Guvenel et al., 2020) have already detected P-specific T cell responses in peripheral blood of healthy adults under experimental RSV inoculation.

In summary, our study provides a new concept for RSV vaccine development, taking into account a balanced Th1/Th17 CD4^+^ T cell response for evaluation of an RSV vaccine. This concept is exemplified by P-KFD1 i.n. immunization that generated RSV P-specific CD4^+^ T cell responses and conferred protection against RSV infection and disease in mice. Overall, the *E. coli*-produced recombinant flagellin-phosphoprotein can prevent RSV infection without raising any VED concerns.

## ACKNOWLEDGMENTS

We thank Professor Zishu Pan at Wuhan University for providing RSV strain A2, and Professor Xu Yang at Central China Normal University for technical assistance in detecting airway responsiveness of mice. We thank Xuefang An, Li Li, He Zhao, Dr. Ding Gao and Juan Min at the core facility and technical support of Wuhan Institute of Virology, CAS, for their kind help in animal experiments and flow cytometry analysis. We are grateful to Yan Wang for assistance in cell sorting by FACS at Institute of Hydrobiology, CAS. We are also grateful to Dr. Yuan Liu and Rulei Yin at Beijing Emei Tongde Technology Development Co., Ltd for their generous support in single cell RNA sequencing and transcriptome analysis. This work was supported by the National Natural Science Foundation of China (No. 31970878) and the “One-Three-Five” Strategic Planning Program of Wuhan Institute of Virology, CAS (No. Y206515SA1).

## AUTHOR CONTRIBUTIONS

B. Z. and J. Y. designed and conducted the experiments, analyzed the data and drafted the manuscript. B. H. and S. L. provided assistance in sc-RNA seq data analysis. X. L., H. Y., Y. Y., B. L., F. Z., X. F. and Y. Z. provided assistance in animal experiments. D. Z., Y. C., M. Z. and E. Z. contributed to discussion and designment of animal studies. S. J. and H. Yan conceived and supervised the work, designed the experiments, analyzed the data, wrote and revised the manuscript.

## STAR Methods

### RESOURCE AVAILABILITY

#### Lead contact

Further information and requests for resources and reagents should be directed to and will be fulfilled by the Lead Contact, Huimin Yan

#### Materials Availability

This study did not generate new unique reagents.

#### Data and Code Availability

### EXPERIMENTAL MODEL AND SUBJECT DETAILS

#### Mice

Female 6-8-week-old BALB/c mice were purchased from Beijing Vital River Laboratory Animal Technology Co. Ltd, Beijing, China. pIgR^−/−^ mice on a BALB/c background were obtained from the Mutant Mouse Resource and Research Centers (MMRRCs), bred and housed at the Animal Center of Wuhan Institute of Virology (WIV), Chinese Academy of Science (CAS). Mice were randomly assigned to groups. All mice were raised in individually ventilated cages (IVCs) under specific pathogen-free (SPF) conditions. The infection experiments were performed in the Animal Biosafety Level 2 (ABSL-2) Laboratory at WIV, CAS. Animal studies were approved by the Animal Welfare and Ethical Review Committee of WIV, and conducted according to Regulations for the Administration of Affairs Concerning Experimental Animals in China (study numbers WIVA09201505 and WIVA09201901).

#### Cell lines

HEp-2 cells were obtained from the China Center for Type Culture Collection (CCTCC), Vero C1008 cells (CRL-1587) and Caco-2 cells (HTB-37) were obtained from the American type culture collection (ATCC). All three cell lines were grown in Dulbecco’s modified Eagle’s (DMEM) medium, supplemented with 10% (v/v) FBS and 1% (v/v) Penicillin/Streptomycin at 37℃ in 5% CO_2_.

#### Virus

Human respiratory syncytial virus (hRSV) strain A2 provided by Professor Pan (Wuhan University) was propagated in HEp-2 or Vero cells.

#### Vaccination

Mice were intranasally (i.n.) immunized three times with P-KFD1 or P+CTB at four-week intervals after anesthesia with pentobarbital sodium (50 mg/kg), and were challenged with 2 ⅹ 10^6^ PFU of RSV A2 on 28 days post last immunization.

Mice were intramuscularly (i.m.) immunized with FI-RSV in the lower hind limb twice at a two-week interval and were challenged with RSV A2 14 days post boost immunization.

#### CFSE labeling and adoptive transfer model

Donor mice were immunized with saline (i.n. or i.m.), KFD1 (i.n.), P-KFD1 (i.n.), P+CTB (i.n.) or FI-RSV (i.m.) as described above. Splenocytes were separated from donor mice at 7 days post second boost immunization and were labeled with 10 μM of CFSE for 10 minutes at room temperature. After washing, CFSE-labeled splenocytes were resuspended in PBS and intravenously injected into naïve mice. Twelve hours post transfer, recipient mice were infected with RSV A2 and sacrificed at 4 days post infection (dpi). Pulmonary and splenic lymphocytes of recipient mice were harvested on 4 dpi for proliferation and Th-bias assay.

### METHOD DETAILS

#### Vaccine preparation

RSV *P* gene was connected to the 5’ flanking region of the *D0-D1* genes of flagellin KF (*E.coli* K12 strain MG1655) to construct the *P-KFD1* gene, which was then cloned into pET-30a plasmid vector (Invitrogen). The recombinant RSV P protein, RSV N protein, HIV-1 p24 protein and P-KFD1 protein were induced, expressed in *E.coli* BL21 DE3, purified by affinity chromatography on a Ni-NTA column (Qiagen), and removed contaminating lipopolysaccharide (LPS), respectively, as previously described (Yang et al., 2013). The residual LPS content was determined using the Limulus assay (Associates of Cape Cod) to be less than 0.01 EU/μg protein.

FI-RSV vaccine was prepared as described elsewhere (Kim et al., 1969; Waris et al., 1996). Briefly, clarified RSV-infected HEp-2 cell culture supernatant was incubated with neutral formalin (1:4000) for 72 hours at 37℃, and the resulting pellet was resuspended and precipitated with Imject Alum adjuvant (Thermo Fisher Scientific) before use.

#### RSV preparation and inoculation

RSV-infected HEp-2 cells were sonicated and concentrated by ultracentrifugation (BECKMAN, Optima XPN) at 70,000 ⅹ *g* for 2 hours at 4℃. The resultant precipitate was resuspended with DMEM. After centrifugation at 10,000 ⅹ *g* for 10 minutes at 4℃, the obtained supernatant was stored at −80℃ as RSV stock. Mice were intranasally inoculated with 2 ⅹ 10^6^ PFU of RSV A2 in 100 μL under avertin (250 mg/kg) anesthesia.

#### Tissue collection and virus titration

RSV-infected mice were sacrificed, noses and the left lungs were removed and placed in a 2-mL screw cap tube containing sterile 1-mm diameter ceramic beads and 1 mL Hanks balanced salt solution (HBSS) containing 25 mM N-2-hydroxyethylpiperazine-N’-2-ethanesulfonic acid (pH 7.8), 200 U of penicillin and streptomycin, 0.218 M sucrose, 4.8 mM glutamate, and 30 mM magnesium chloride (Taylor et al., 1984; Waris et al., 1996). The tubes were quick-frozen in liquid nitrogen. Then the tissue homogenates were prepared using a BeadBeater homogenizer (Biospec Products) for 4 minutes with intermittent cooling in ice bath (Waris et al., 1996). Virus stocks or tissue homogenates from infected mice were titrated by immuno-plaque assay as described elsewhere (Quan et al., 2011). Titers were recorded as PFU/nose or PFU/g lung.

#### Enzyme-Linked Immunosorbent Assay (ELISA)

Antibody responses were assessed by enzyme-linked immunosorbent assay (ELISA). In brief, a 96-well plate was coated with target protein (3 μg/mL) in carbonate-bicarbonate buffer at 4℃ overnight and then blocked with 1% BSA at 37℃ for 2 hours. Then serial four-fold diluted samples were added to the plates for 2 hours at 37℃. After washing, secondary alkaline phosphatase-labeled antibodies (SouthernBiotech) were applied to the plates followed by substrate (p-nitrophenyl phosphate, Sigma) coloring. ODs were read at 405 nm by an ELISA plate reader (Thermo Labsystems).

IFN-γ, IL-4, IL-17A and IL-8 production in cultured cell supernatants was detected by ELISA kits (eBioscience) under the manufacturer’s instructions.

#### Airway responses to methacholine challenge

Airway responsiveness was assessed in mice using an AniRes2005 lung function system (Bestlab, version 2.0, China) according to the manufacturer’s instructions, as described previously (Drazen et al., 1999; Guo et al., 2012). Briefly, anesthetized mice were ventilated via a tracheal cannula which was connected to an animal ventilator, then methacholine was administered through a catheter into the jugular vein with increasing concentrations (0.025, 0.05, 0.1 and 0.2 mg/kg) at 5-minute intervals. The relative area (R-area) was defined as the area between the peak value and baseline of Ri or Re during a 250-second period, and the valley value of Cdyn was used to describe lung compliance.

#### Histology

Lungs of mice were excised and fixed in 10% neutral buffered formalin for 24 hours at room temperature, followed by embedding in paraffin. After cutting into slices, tissues were stained with hematoxylin & eosin (H&E) or periodic acid-Schiff (PAS). Whole slide imaging was observed using the slide scanner Pannoramic MIDI (3DHISTECH). Pathological changes were evaluated as described elsewhere and scored on a 1-4 severity scale (Knudson et al., 2015). Scores of inflammatory cell aggregation and interstitial pneumonia scale: 1-normal naïve parameters, 2 -slight and occasional cell aggregation, 3-moderate cell infiltration, 4-moderate to severe and multifocal cell aggregation around bronchioles or pulmonary vessels or air space of lung sections. Scores of mucus production scale: 1-no mucus detectable, 2-rare mucus, 3-moderate mucus accumulation, 4-severe mucus production in airways.

#### Isolation of immunocytes from tissues

For isolation of pulmonary immunocytes in mice, lungs were removed and cut into pieces and digested in 2 mL of HBSS buffer (HyClone) containing 0.5 mg/mL collagenase 1 (Gibco) and 0.2 mg/mL Dnase 1 (Sigma) for 45 minutes at 37℃. After digestion, lung pieces were pressed through a 70-μm nylon mesh screen, and the resulting single-cell suspensions were collected and washed with cold phosphate buffer saline (PBS). Then the cell pellets were resuspended in 40% Percoll (GE Healthcare) and layered carefully onto 70% Percoll to generate discontinuous Percoll gradients and centrifuged at 2000 rpm for 25 minutes at 22℃. Cells were aspirated in the interface between 40% and 70% Percoll gradients.

For isolation of splenic lymphocytes in mice, spleens were minced and pressed through the 70-μm nylon mesh screen. Lymphocytes were obtained by density centrifugation described above using Mouse Lymphocytes Separation Medium (Dakewe).

#### Flow cytometry

To detect the number of eosinophils and T cells in lung, single-cell suspensions from lungs were incubated with anti-CD16/CD32 (93) to block nonspecific Fc receptor binding. Then cells were surface-stained with monoclonal antibodies specific to CD45 (30-F11), CD11b (M1/70), Siglec F (E50-2440), Gr-1 (RB6-8C5), CD4 (RM4-5), CD8 (53-6.7) and fixable viability dye (eBioscience) or 7-AAD viability staining solution (BioLegend) at 4℃ for 30 minutes. For Th bias assay, separated cells were stimulated for 5 hours at 37℃ with 200 ng/mL PMA (Beyotime) and 2000 ng/mL ionomycin (Beyotime) in the presence of brefeldin A (BioLegend) and monensin (BioLegend). Then cells were labeled with fixable viability dye and mAbs specific to CD4 or CD8. After washing, cells were fixed and permeabilized, using the fixation buffer and Intracellular Staining Permeabilization Wash Buffer (BioLegend), and stained intracellularly with mAbs specific to IFN-γ (XMG1.2), IL-4 (11B11) and IL-17A (eBio17B7). For detection of Treg, cells were labeled with 7-AAD viability staining solution and surfaced-stained with mAbs specific to CD4 and CD25 (PC61) and then fixed and permeabilized with Foxp3/Transcription Factor Staining Buffer Set (eBioscience) prior to incubation with mAb specific to Foxp3 (FJK-16s). For detection of cell proliferation, pulmonary and splenic lymphocytes of recipient mice were separated and stained with fixable viability dye and mAbs specific to CD4, CD8, and B220 (RA3-6B2) to analyze the expanded CFSE-labeled CD4^+^ T cells, CD8^+^ T cells and B cells by flow cytometry. All antibodies were bought from BD Bioscience, eBioscience or BioLegend. Stained cells were run on an LSRFortessa (BD, Heidelberg, Germany) and analyzed with FlowJo software (Tree Star, Ashland, OR).

#### Enzyme-Linked Immunospot (ELISpot) Assay

Mouse IFN-γ (Dakewe), mouse IL-17A (Mabtech) and mouse IL-4 (Dakewe) secreting cells were detected by ELISpot following the manufacturer’s instructions. Briefly, 2×10^5^ of lymphocytes were seeded into precoated ELISpot plates and stimulated with target proteins, or irrelevant protein (HIV-1 p24), for 45 hours at 37℃, 5% CO_2_, respectively. After cell lysis, biotin-conjugated mAbs were applied prior to adding streptavidin-HRP, and then spots were developed with an AEC coloring system. The number of spots was counted by an automatic ELISpot reader (AID, Germany).

#### Serum transfer

Serum from saline, FI-RSV (i.m.) or P-KFD1 (i.n.) immunized mice were collected at 14 days post last immunization and inactivated at 56℃ for 30 minutes. Then, 250 μL of filtered serum were injected into naïve recipient (per mouse), respectively, followed by RSV infection.

#### *In vivo* antibody treatment

For depletion of CD4^+^, CD8^+^, or both CD4^+^ and CD8^+^ T cells, mice were intraperitoneally injected with 250 μg of α-CD4 antibody (GK1.5), 250 μg of α-CD8 antibody (2.43) or 250 μg of α-CD4 antibody plus 250 μg of α-CD8 antibody two days before RSV infection, respectively. Depletion antibodies were injected twice at a four-day interval (Knudson et al., 2015). Mice treated with the same doses of isotype-matched rat IgG2b (LTF-2) antibody were designed as controls.

For blocking IFN-γ, IL-17A or both IFN-γ and IL-17A, mice were intraperitoneally injected with 250 μg of α- IFN-γ antibody (XMG1.2), 250 μg of α-IL-17A antibody (17F3), or 250 μg of α-IFN-γ antibody, together with 250 μg of α-IL-17A antibody, respectively. Blocking antibodies were injected every other day starting 1 day before RSV challenge. Mice treated with the same doses of rat IgG 1 (HRPN), mouse IgG 1 (MOPC-21), or both rat IgG 1 and mouse IgG 1 were assigned as isotype control groups, respectively.

All antibodies used above were purchased from Bio X Cell.

#### FTY720 treatment

Mice were intraperitoneally injected with 250 μL of FTY720 (Sigma) at a dose of 1 mg/kg daily beginning 3 days before RSV challenge, until they were sacrificed.

#### Purification of splenic CD4^+^ T cells and adoptive transfer

Splenocytes were separated from saline or P-KFD1 immunized mice at 7 days post last immunization. Then CD4^+^ T cells were positively selected using anti-CD4 (L3T4) microbeads following the manufacturer’s instructions (Miltenyi Biotec, Germany). The purity of sorted CD4^+^ T cells was at least 95%. Naïve recipient mice were intravenously injected with 3×10^6^ purified CD4^+^ T cells 12 hours prior to RSV challenge and sacrificed at 4 days post infection.

#### FACS sorting, co-culture and sc-RNA seq

Firstly, CD4^+^ T cells were positively isolated by magnetic sorting (Miltenyi Biotec, Germany) from pulmonary immunocytes of saline or P-KFD1 intranasally immunized mice on 4 dpi. Then the purified CD4^+^ T cells were stained with monoclonal antibodies specific to CD3 (clone 145-2C11), CD4 (clone RM4-5), CD11a (clone M17/4), CD49d (clone 9C10) and 7-AAD viability staining solution (BioLegend) for FACS sorting on a FACSAriaIII (BD, Heidelberg, Germany).

For detection of P-specific immune responses, the FACS sorted CD11a^high^ CD4^+^ T cells and CD11a^low^ CD49d^−^ CD4^+^ T cells were stimulated with RSV P protein or irrelevant protein or medium alone in 96-well plates for 48 hours, respectively. In detail, 3ⅹ10^4^ sorted cells were co-cultured with 1ⅹ10^4^ naïve mice derived splenocytes in a well. And the collected cell supernatants were used to detect the amount of IFN-γ, IL-4 and IL-17A by ELISA.

For sc-RNA seq, FACS sorted CD11a^low^ CD49d^−^ CD4^+^ T cells were labelled by TotalSeq™ anti-mouse Hashtag antibody (BioLegend) and then mixed with the corresponding CD11a^high^ CD4^+^ T cells at a ratio approximately 3:5. Sc-RNA seq library preparation was conducted under the manufacturer’s instructions for the 10x Genomics 5’ v1.0 chemistry with immune profiling and feature barcoding technology for cell surface protein. Briefly, 14000 cells were loaded on the chip which targeted 8000 cells corresponding to the manufacturer’s guidance. A single cell, a barcoded gel bead, and reverse transcriptase reagent were encapsulated into a Gel-Bead-In-EMulsion (GEM). GEMs were transferred to the 8-tube strip and incubated at 53℃, 45 min followed by 85℃, 5 min for reverse transcription and enzyme inactivation respectively. Initial amplification of cDNA and library preparation were carried out with 13 and 14 cycles of amplification, respectively; V(D)J libraries were generated corresponding to each 5’ gene expression library using both 10 cycles for the two enrichment PCRs and 10 cycles for the library amplification, respectively (Ni et al., 2020). TotalSeq™-A Hashtag antibodies which were not compatible with the standard cell surface protein library procedure were used in this study. Therefore, the Hashtag cell surface libraries were constructed as following steps. Partial of the amplified cDNAs were ligated with illumina read 2 adaptor, then with the indexing PCR to get the Hashtag libraries. Libraries were sequenced on Illumina NovaSeq6000 sequencing platform with the following read lengths: read 1-150 cycles; read 2-150 cycles; and i7 index-8 cycles.

#### Single cell transcriptome analysis

The Cell Ranger Single-Cell Software Suite (versions 3.0.2) were used to perform barcode processing and single-cell gene counting (Paulson et al., 2018) (http://10xgenomics.com/). First, “cellranger mkfastq” was carried to generated fastq files. Second, feature-barcode matrices were generated by “cellranger count” function using GRCm38 mouse as reference genome (Ensembl). Then the cell ranger aggregation function (aggr) was used to combine the two libraries. A correction for sequencing depth was also performed during the aggregation. IntegrateData function in R package Seurat V3.6.3 was used to merge Saline and P-KFD1 fastq data. Low-quality genes and cells were filtered by removing cells with 1) expressed genes fewer than 200, 2) expressed genes more than 5,000, 3) percentages of mitochondrial genes >20% and 4) genes expressed in less than 3 cells. The filtered gene-barcode matrix was first normalized using “LogNormalize” method. The top 2,000 variable genes were then identified using the “vst” method in Seurat FindVariableFeatures function. Principal component analysis (PCA) was performed using variable genes, and the top 20 principal components (PCs) were used to perform t-distributed Stochastic Neighbor Embedding (t-SNE) to visualize the cells. The resolution was set to 0.5 for clustering. T-SNE coordinate points and cell clusters were exported into Loupe Cell Brower V3.1.1 (combine data by cell ranger aggregation function) to analyze data. For the TCR data, the Cell Ranger Single-Cell Software Suite (versions 3.0.2) were used to perform barcode processing, assembly contig, cell calling, annotation the contigs and CDR3 region for each clonotype. KEGG (Kyoto Encyclopedia of Genes and Genomes) pathways and GO (Gene ontology) terms analysis were performed as follows: genes with Benjamini-Hochberg-adjusted P value < 0.05 and log_2_|FC| between two groups larger than 1 were used for DAVID analysis (Guo et al., 2018) (https://david-d.ncifcrf.gov/).

### QUANTIFICATION AND STATISTICAL ANALYSIS

Statistical parameters including the exact value of n, the definition of center, dispersion, and precision measures (geometric mean ± SEM) and statistical significance are displayed in Figures and Figure Legends. Data were considered to be statistically significant if p < 0.05. All statistical analyses were performed using GraphPad Prism version 5.0 (GraphPad Software, San Diego, CA). Data were compared using unpaired, two-tailed *t* test between two groups or one-way ANOVA with Tukey’s multiple comparison for more than two groups. Data were also analyzed by regular two-way ANOVA if two independent variables existed in one experiment.

