## supplemental figures and figure legends for "A Safe and Effective Mucosal RSV Vaccine Consisting of RSV Phosphoprotein and Flagellin Variant"

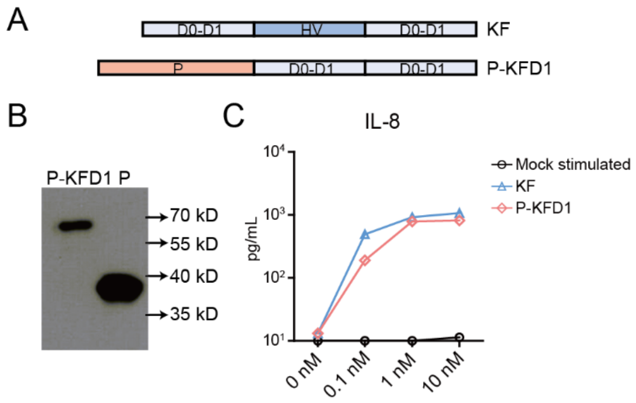


**Figure S1. Recombinant protein P-KFD1 retains TLR5 agonist activity, Related to Figure 1.** (**A**) Schematic diagram of fusion protein P-KFD1. *P* gene was connected to the 5’ flanking region of the *D0* and *D1* genes derived from flagellin KF to generate *P-KFD1*. (**B**) Western blot detection of P and P-KFD1 protein with P-specific monoclonal antibody. (**C**) IL-8 production in supernatants of cultured epithelial cell line Caco-2 cells with KF or P-KFD1 stimulation was detected by ELISA, n = 3 replicates per group. Data are represented as mean ± SEM of three independent experiments.

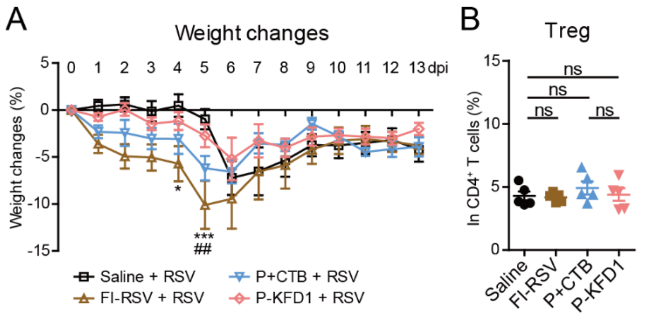


**Figure S2. Weight changes and percentage of Tregs in lungs of immunized mice after RSV challenge, Related to Figure 2.** (**A**) Weight changes in saline, FI-RSV, P+CTB or P-KFD1 immunized mice post RSV challenge, n = 6 mice per group. (**B**) Percentages of $CD25^+$ $Foxp3^+$ Tregs in $CD4^+$ T cells in lungs of immunized mice at 8 dpi, n = 5 mice per group. Data are represented as mean ± SEM of two independent experiments. In **Figure S2A**, groups were compared using regular two-way ANOVA. * $p < 0.05$, *** $p < 0.001$, significant difference between FI-RSV group and saline group. ## $p < 0.01$, significant difference between FI-RSV group and P-KFD1 group. In **Figure S2B**, groups were compared using one-way ANOVA, ns means non-significant.

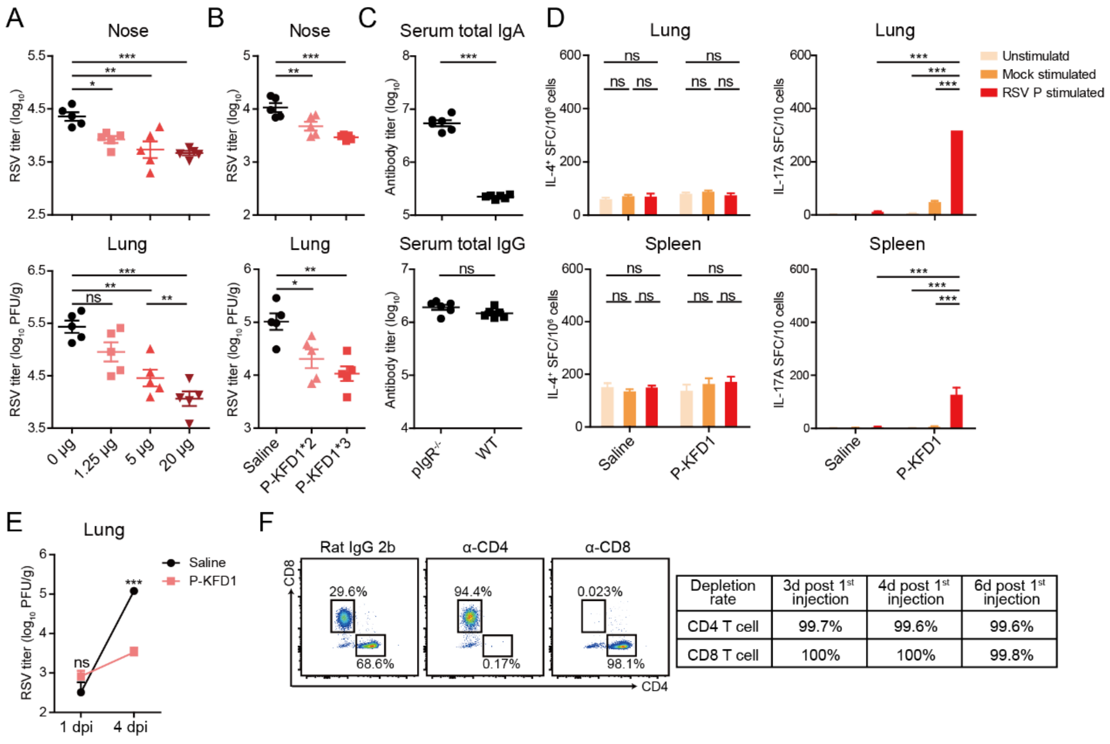


| Depletion rate | 3d post 1st injection | 4d post 1st injection | 6d post 1st injection |
|---|---|---|---|
| CD4 T cell | 99.7% | 99.6% | 99.6% |
| CD8 T cell | 100% | 100% | 99.8% |

**Figure S3. Protection associated factors in P-KFD1 intranasally immunized mice, Related to Figure 3.** (**A**) Groups of mice were intranasally immunized with 0, 1.25, 5 or 20 μg of P-KFD1 three times followed by RSV challenge, respectively, and viral loads in noses and lungs of mice were detected at 4 dpi, n = 5 mice per group. (**B**) Groups of mice were intranasally immunized with 10 μg of P-KFD1 twice or three times prior to RSV challenge, respectively, and viral loads in noses and lungs of mice were detected at 4 dpi, n = 5 mice per group. (**C**) Serum total IgA and IgG antibody titers in pIgR$^{-/-}$ and wild type mice were detected by ELISA, n = 6 mice per group. (**D**) P-specific IL-4$^{+}$ T cell or IL-17A$^{+}$ T cell responses in lungs and spleens of saline or P-KFD1 immunized mice were detected by ELISpot on 7 days post last immunization, n = 6 mice per group. (**E**) Mice were intranasally immunized with saline or P-KFD1, prior to RSV challenge, respectively, RSV titers were monitored at 1 dpi and 4 dpi, n = 5 mice per group. (**F**) Mice were intraperitoneally injected with 250 μg of anti-CD4 or anti-CD8 antibodies on day 0 and day 4, respectively, n = 2 mice per treatment. Blood from treated mice was collected on day 3, day 4 and day 6, and the frequencies of $CD4^+$ or $CD8^+$ T cells were detected by FCM. Representative plots and cumulative results of depletion rates in mice that were treated with anti-CD4 or anti-CD8 antibodies were displayed. In **Figures S3A-S3E**, data are represented as mean ± SEM of at least two independent experiments. In **Figures S3A and S3B**, groups were compared using one-way ANOVA. In **Figure S3C**, two groups were compared using unpaired *t* test. In **Figures S3D and S3E**, groups were compared using regular two-way ANOVA. * $p < 0.05$, ** $p < 0.01$, *** $p < 0.001$, ns means non-significant.

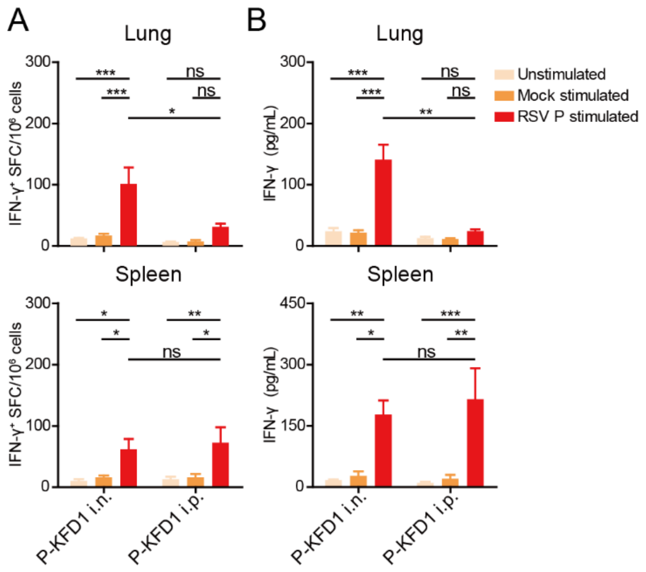


**Figure S4. Intranasal or intraperitoneal immunization with P-KFD1 induced P-specific IFN-γ$^{+}$ T cell response, Related to Figure 4.** Groups of mice were intranasally or intraperitoneally immunized with P-KFD1 three times at 4-week intervals and sacrificed at 7 days post last immunization. (**A**) P-specific IFN-γ secreting T cells in lungs and spleens of immunized mice were tested by ELISpot, n = 5 mice per group. (**B**) Lymphocytes from lungs or spleens of immunized mice were incubated with P protein overnight, and IFN-γ production in supernatants was determined by ELISA, n = 5 mice per group. Data are represented as mean ± SEM of two independent experiments. Groups were compared using regular two-way ANOVA. * $p < 0.05$, ** $p < 0.01$, *** $p < 0.001$, ns means non-significant.

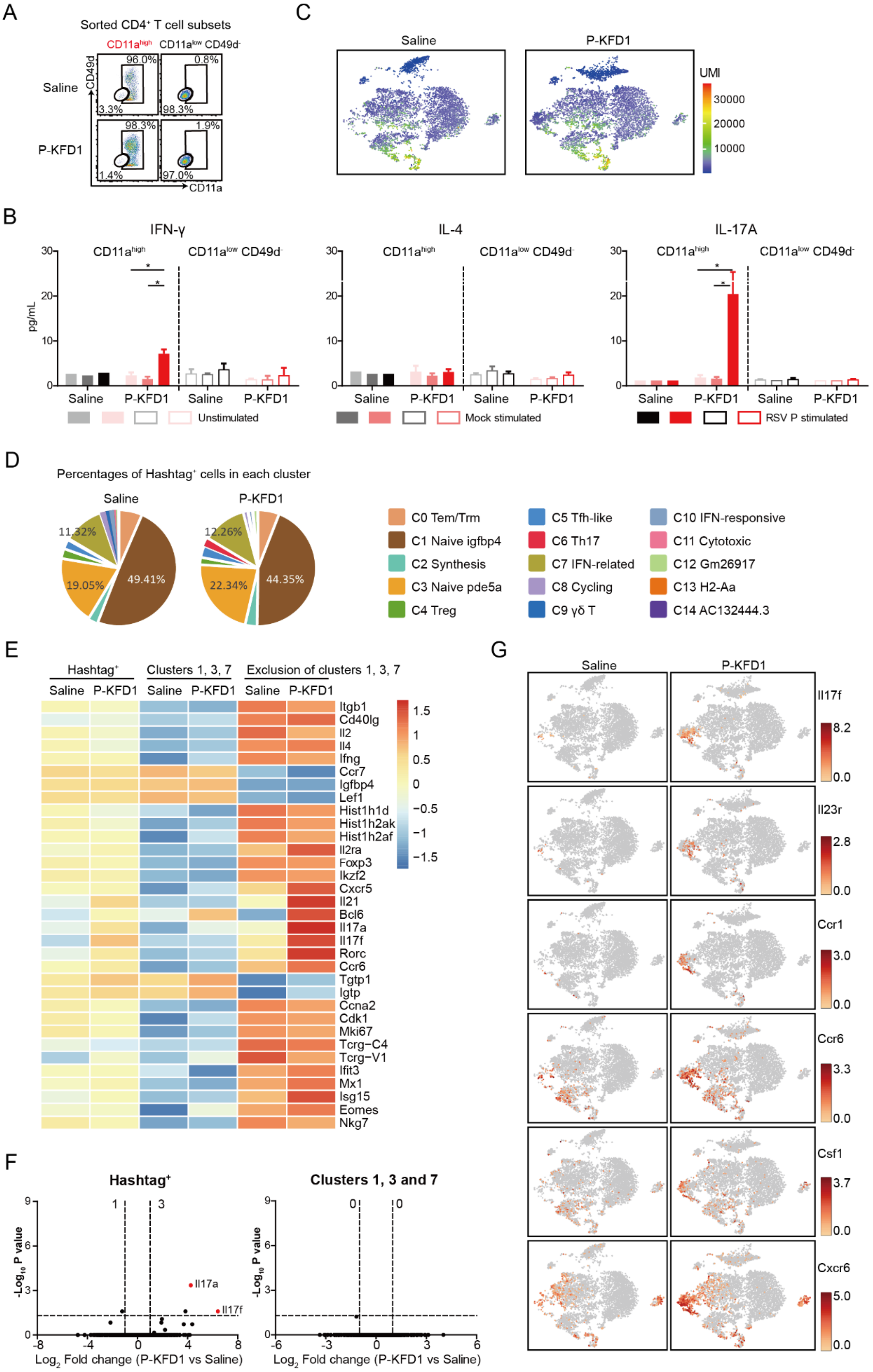
A
Sorted CD4+ T cell subsets
CD11a high
CD11a low CD49d-
Saline
P-KFD1
CD49d
CD11a
96.0%
0.8%
3.3%
98.3%
98.3%
1.9%
1.4%
97.0%
C
Saline
P-KFD1
UMI
30000
20000
10000
B
IFN-γ
IL-4
IL-17A
pg/mL
CD11a high
CD11a low CD49d-
Saline
P-KFD1
Unstimulated
Mock stimulated
RSV P stimulated
D
Percentages of Hashtag+ cells in each cluster
Saline
P-KFD1
11.32%
49.41%
19.05%
12.26%
22.34%
44.35%
C0 Tem/Trm
C1 Naive igfbp4
C2 Synthesis
C3 Naive pde5a
C4 Treg
C5 Tfh-like
C6 Th17
C7 IFN-related
C8 Cycling
C9 γδ T
C10 IFN-responsive
C11 Cytotoxic
C12 Gm26917
C13 H2-Aa
C14 AC132444.3
E
Hashtag+
Clusters 1, 3, 7
Exclusion of clusters 1, 3, 7
Saline
P-KFD1
Itgb1
Cd40lg
Il2
Il4
Ifng
Ccr7
Igfbp4
Lef1
Hist1h1d
Hist1h2ak
Hist1h2af
Il2ra
Foxp3
Ikzf2
Cxcr5
Il21
Bcl6
Il17a
Il17f
Rorc
Ccr6
Tgtp1
Igtp
Ccna2
Cdk1
Mki67
Tcrg−C4
Tcrg−V1
Ifit3
Mx1
Isg15
Eomes
Nkg7
1.5
1
0.5
0
−0.5
−1
−1.5
F
Hashtag+
Clusters 1, 3 and 7
-Log10 P value
Log2 Fold change (P-KFD1 vs Saline)
Il17a
Il17f
G
Saline
P-KFD1
Il17f
8.2
0.0
Il23r
2.8
0.0
Ccr1
3.0
0.0
Ccr6
3.3
0.0
Csf1
3.7
0.0
Cxcr6
5.0
0.0

**Figure S5. In CD4$^+$ T cells, P-KFD1 i.n. immunization mainly changes the antigen-experienced CD11a$^{high}$ and non-naive CD4$^+$ T cells, Related to Figure 5.** (**A**) Purity of FACS sorted CD11a$^{high}$ CD4$^+$ and CD11a$^{low}$ CD49d$^-$ CD4$^+$ T cell subsets from saline or P-KFD1 intranasally immunized mice on 4 dpi. (**B**) IFN-γ, IL-4 and IL-17A production in FACS sorted CD11a$^{high}$ CD4$^+$ or CD11a$^{low}$ CD49d$^-$ CD4$^+$ T cells post stimulation. FACS sorted cells were cultured on 96-well plates and stimulated with P protein or irrelevant protein or medium for 48 hours, respectively. (**C-G**) Sc-RNA sequencing analysis. (**C**) Distribution of unique molecular identifiers (UMI) in each cluster on the t-SNE plots of saline and P-KFD1 groups. (**D**) Pie charts for the proportions of hashtag$^+$ cells in each cluster. (**E**) Heatmap of clustering identification associated DEGs in hashtag$^+$ cells, clusters 1, 3, 7 cells, and exclusion of clusters 1, 3, 7 cells. (**F**) Volcano plots of DEGs in hashtag$^+$ cells or in clusters 1, 3, 7 cells between P-KFD1 group versus saline group. (**G**) T-SNE plots for gene expression of *Il17f, Il23r, Ccr1, Ccr6, Csf1* and *Cxcr6*. Data are represented as mean ± SEM of two independent experiments for **Figure S5B**. Groups were compared using regular two-way ANOVA. * $p < 0.05$.

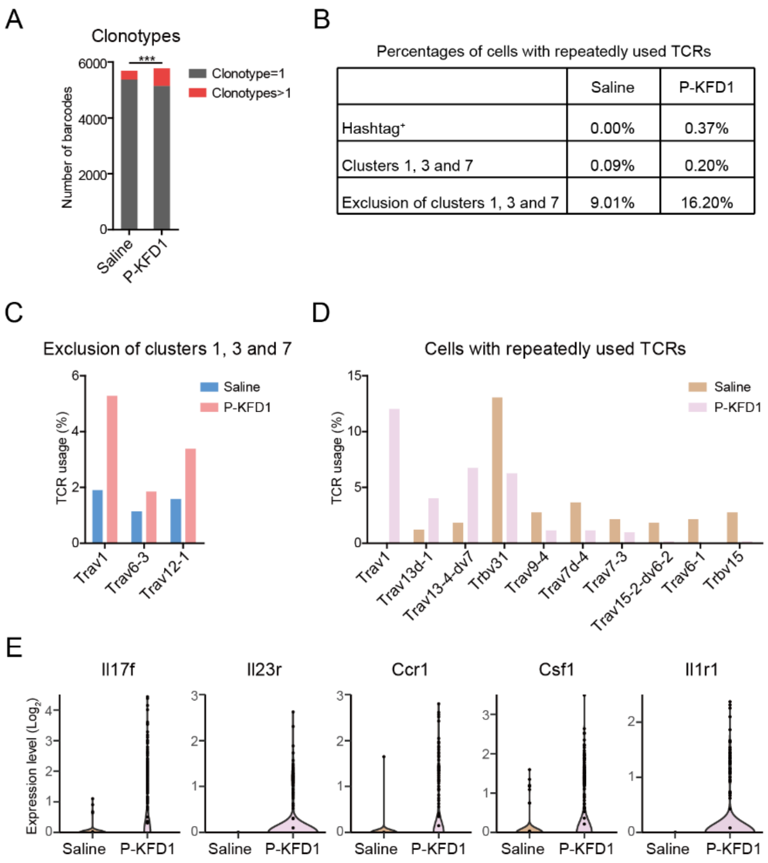


**Figure S6. Clonal expansion of CD4$^+$ T cells in saline or P-KFD1 intranasally immunized mice on 4 dpi, Related to Figure 6.** (**A**) Number of CD4$^+$ T cells possessed a single clonotype or with repeatedly used TCRs. The difference in cells with repeatedly used TCRs or single used TCRs between saline group and P-KFD1 group were calculated by chi-square test. *** $p < 0.001$. (**B**) Percentages of cells with repeatedly used TCRs in hashtag$^+$ cells, clusters 1, 3, 7 cells or exclusion of clusters 1, 3, 7 cells in saline and P-KFD1 group. (**C** and **D**) The difference of alpha and beta chain of TCR usage in cells excluded clusters 1, 3 and 7 (**C**) and in cells with repeatedly used TCRs (**D**) between saline and P-KFD1 group. (**E**) Violin plots for gene expression of *Il17f*, *Il23r*, *Ccr1*, *Csf1* and *Il1r1* in cells with repeatedly used TCRs.

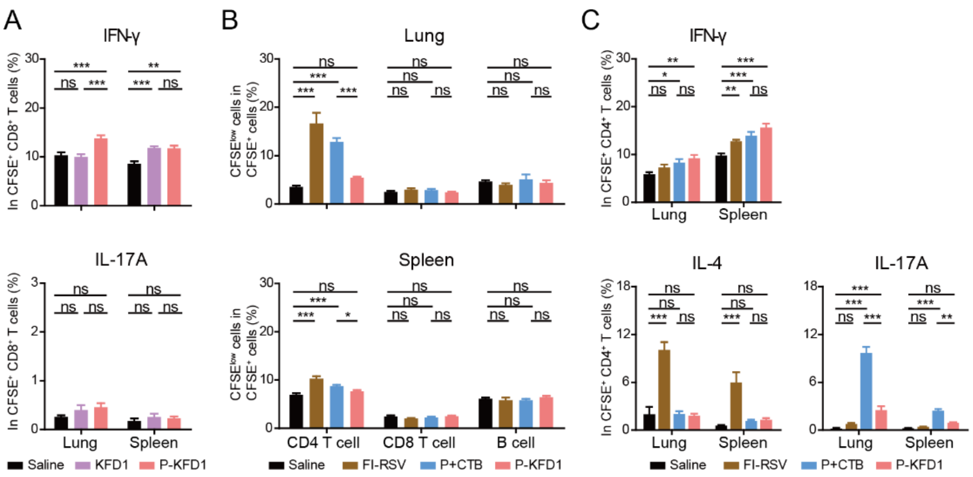


**Figure S7. Intranasal immunization with P-KFD1 elevated IFN-γ$^+$ T cell and IL-17A$^+$ T cell responses, Related to Figure 7.** (**A-C**) Adoptive transfer experiments. (**A**) Recipient mice received CFSE-labeled splenocytes from saline, KFD1 or P-KFD1 intranasally immunized mice, respectively, followed by RSV challenge. With PMA and ionomycin stimulation, percentages of IFN-γ$^+$ cells or IL-17A$^+$ cells in CFSE$^+$ CD8$^+$ T cells in lungs and spleens of recipient mice were measured at 4 dpi, n = 4~5 mice per group. (**B-C**) Recipient mice received CFSE-labeled splenocytes from saline, FI-RSV, P+CTB or P-KFD1 immunized donor mice, respectively, followed by RSV challenge. (**B**) Percentages of proliferated CD4$^+$, CD8$^+$ T cells and B cells in CFSE$^+$ cells in lungs and spleens of recipients were evaluated at 4 dpi, n = 4~5 mice per group. (**C**) Percentages of IFN-γ$^+$, IL-4$^+$ or IL-17A$^+$ cells in CFSE$^+$ CD4$^+$ T cells in lungs and spleens of recipients were assessed at 4 dpi, n = 4~5 mice per group. Data are represented as mean ± SEM of two independent experiments. In **Figures S5A and S5C**, groups were compared using one-way ANOVA. In **Figure S5B**, groups were compared using regular two-way ANOVA. * $p < 0.05$, ** $p < 0.01$, *** $p < 0.001$, ns means non-significant.
